# A *Plasmodium falciparum* PX1 haplotype is associated with reduced susceptibility to artemisinin and lumefantrine

**DOI:** 10.64898/2026.05.05.722990

**Authors:** Christopher Bower-Lepts, Arvind Jangra, Sachie Kanatani, Abhai Tripathi, Godfree Mlambo, Alejandro McCotter-González, Linda Romano, Stephen Orena, Martin Okitwi, Karamoko Niaré, Philip J. Rosenthal, Melissa Conrad, Photini Sinnis, Sachel Mok

**Affiliations:** Division of Infectious Diseases, Department of Medicine, Columbia University Irving Medical Centre, New York, NY, USA; Centre for Malaria Therapeutics and Antimicrobial Resistance, Columbia University Irving Medical Centre, New York, NY, USA; Aaron Diamond AIDS Research Centre (ADARC), Columbia University Irving Medical Centre, New York, NY, USA; Department of Molecular Microbiology & Immunology and the Johns Hopkins Malaria Research Institute, Johns Hopkins School of Public Health, Baltimore, MD, USA; Infectious Diseases Research Collaboration, Kampala, Uganda; Department of Pathology and Laboratory Medicine, Brown University, Providence, RI, USA; Pathogens Genomics Diversity Network Africa, Imm. Gwancoura, Sotuba, Bamako, Mali; Department of Medicine, University of California, San Francisco, CA, USA; Division of Infectious Diseases, Department of Medicine, Johns Hopkins School of Medicine, Baltimore MD, USA

## Abstract

Effective control of falciparum malaria depends on the sustained efficacy of frontline antimalarial drugs, particularly artemether-lumefantrine (AL), the most widely used therapy in Africa. However, the emergence of artemisinin partial resistance and reduced lumefantrine susceptibility in eastern Africa threaten malaria control and elimination. Robust genetic markers of decreased susceptibility to lumefantrine remain elusive, and our understanding of artemisinin resistance is incomplete. We report results of a *Plasmodium falciparum* genetic cross between a drug-sensitive line and a Ugandan strain exhibiting reduced susceptibility to dihydroartemisinin and lumefantrine. Targeted deep sequencing of progeny pools and 460 recombinant progeny clones derived under drug pressures revealed distinct haplotypic signatures. Drug-selection experiments identified genetic polymorphisms in *Plasmodium falciparum px1*, encoding a phosphoinositide-binding protein, as the strongest correlates of reduced susceptibility to dihydroartemisinin and lumefantrine. The PX1 PIN haplotype (L1222P, M1701I, D1705N) recently discovered in Ugandan parasites was highly enriched following dihydroartemisinin or lumefantrine treatment of pooled mixtures of genetically diverse Ugandan clinical isolates. This haplotype was associated with reduced susceptibility to dihydroartemisinin and lumefantrine, compared to wild-type sequence, in culture-adapted Ugandan *P. falciparum* lines. These results confirm that PX1 mutations were selected across geographically distinct Ugandan parasite backgrounds. Long-term competitive fitness assays demonstrated that PX1 mutations confer asexual blood-stage parasites with a growth advantage, potentially explaining a rapid rise of PX1 PIN alleles over the last two decades in Uganda. Overall, our data suggest the PX1 PIN haplotype is a robust marker of reduced AL susceptibility in African *P. falciparum*, enabling surveillance of emerging drug resistance.

## Introduction

Malaria remains a major global health challenge, causing an estimated 282 million cases and 610,000 deaths in 2024^1^. Over 94% of cases and deaths occur in the WHO African Region, predominantly due to infections with *Plasmodium falciparum*^1^. Malaria control depends critically on effective antimalarial chemotherapy, with artemisinin-based combination therapies (ACTs) the cornerstone of treatment for uncomplicated disease^2,3^. ACTs combine a rapidly acting artemisinin derivative (artesunate, artemether, or dihydroartemisinin (DHA)) that swiftly reduces parasite density with a longer-acting partner drug that eliminates residual parasites and limits the risk of recrudescence^4^. Since the early 2000s, artemether-lumefantrine (AL) has been the frontline ACT across most of sub-Saharan Africa, including Uganda, which bears the third highest global malaria burden^5^. Sustained efficacy of AL is therefore critically important for malaria control in the region.

After earlier gains, progress against malaria has largely stalled in Africa over the last decade, partly due to the emergence and spread of drug resistance. Artemisinin partial resistance (ART-R), defined by delayed parasite clearance after treatment with an artemisinin or increased parasite survival after brief *in vitro* exposure to an artemisinin, first emerged in the mid-2000s in the Greater Mekong Subregion and is largely confined to Southeast Asia^6^. This phenotype is conferred by mutations in the PfKelch13 (K13) propeller domain, which reduce hemoglobin endocytosis, thereby limiting artemisinin activation via heme-dependent processes^7–9^. In the absence of *k13* mutations, variants in additional genes, including *coronin*, *ubp1*, and *ap2mu,* have also been implicated in modulating artemisinin susceptibility, suggesting a pleiotropic mechanism of resistance^10,11^. Although ART-R was initially confined to Southeast Asia, K13 mutations validated as markers of ART-R have independently emerged in distinct sites in East Africa and the Horn of Africa^12^. These include the A675V and C469Y mutations, which have rapidly expanded to over 50% combined prevalence in parts of northern Uganda, the R561H mutation which is prevalent in Rwanda and neighboring regions of Tanzania and Uganda, and the R622I mutation which has been observed in parts of Eritrea and Ethiopia^13–17^.

While AL has largely retained clinical efficacy in Africa, therapeutic efficacy studies have reported AL treatment failure rates exceeding 10% at multiple sites, including Uganda, Angola, Tanzania, Burkina Faso, and Eritrea^18–22^. In parallel, declining *ex vivo* susceptibility to lumefantrine (LUM) has been reported in northern and eastern Uganda^10,23^, although high-level LUM resistance has not clearly been documented^24,25^. Field studies suggest that AL selects for wild-type alleles at loci where mutations are associated with aminoquinoline resistance, including PfMDR1 (N86Y and D1246Y) and PfCRT (K76T)^26^. These wild-type alleles have both been linked to reduced LUM susceptibility, but with only modest phenotypic effects likely insufficient to explain emerging treatment failures^27,28^. Increased *mdr1* copy number has also been linked to decreased susceptibility to LUM and the related aryl aminoalcohol mefloquine (MEF), which is used as an ACT partner drug in combination with artesunate in Southeast Asia^29,30^. However, *mdr1* copy number amplifications are rare in Africa^31^. Identifying robust genetic markers of LUM resistance remains an urgent need.

Whole-genome sequencing of Ugandan clinical isolates has revealed parasite adaptation in the context of heavy use of AL in the country^32^. Recently, Niaré et al (2025) reported selection of a haplotype (L1222P, M1701I, D1705N; PIN) in a phosphoinositide-binding protein (PX1), in Uganda over the last two decades. The PIN haplotype was seen to emerge prior to the emergence of ART-R-mediating K13 mutations in Uganda and to be associated with decreased susceptibility to lumefantrine, mefloquine, and the active metabolite of artemether, DHA^32^. However, independent *in vitro* evidence linking PX1 field mutations to reduced susceptibility to the components of AL has not yet been established, underscoring the need for robust experimental validation of PX1 as a molecular marker of partner-drug resistance.

Existing strategies to identify the determinants of reduced susceptibility to LUM, including *in vitro* resistance selections and other forward genetic screens, have not yielded validated genetic markers^33^. *P. falciparum* genetic crosses have previously proven highly effective for mapping determinants of antimalarial resistance to chloroquine, piperaquine and artemisinins^34–37^. The recent implementation of the FRG-NOD humanized mouse model, which supports *P. falciparum* liver-and blood-stage infection, has overcome dependence on non-human primate models and greatly facilitated the application of genetic crosses in malaria research^38^. Here, we leveraged this platform to implement a genetic cross between a Ugandan clinical isolate with reduced DHA and LUM susceptibility and the drug-sensitive NF54 strain. We combined orthogonal approaches including drug selection, parasite cloning and competitive fitness assays in cross progeny, with analyses of Ugandan clinical isolates, to identify loci associated with reduced susceptibility to DHA and LUM.

## Results

### Implementation of a genetic cross to map determinants of decreased susceptibility to dihydroartemisinin and lumefantrine

To identify genetic determinants of decreased susceptibility to the components of AL, we implemented an experimental genetic cross between the African clinical isolate, PAT-039 (collected in Agago District, northern Uganda, in 2021) and the drug-sensitive laboratory strain NF54 (with origin from Rwanda in the 1970s, prior to the introduction of AL). PAT-039 demonstrated ART-R, with 19% survival in the 4h ring-stage survival assay (RSA) and a 1.6-fold higher DHA IC₅₀ compared with NF54 (Fig. 1a). PAT-039 also showed reduced susceptibility to LUM (IC₅₀ = 127 nM) and MEF (IC₅₀ = 54.4 nM), corresponding to 3.1-fold and 2.6-fold increases in IC₅₀ relative to NF54, respectively (Fig. 1b,c). Full drug susceptibility profiles are shown in Extended Data Fig. 1 and Supplementary Table 1. Sequencing of loci potentially linked to drug resistance demonstrated that PAT-039 harbors the ART-R-linked K13 C469Y mutation, the PX1 PIN haplotype (L1222P, M1701I, D1705N) and the quintuple mutant DHFR (N51I, C59R, S108N) and DHPS (A437G, K540E) haplotype that mediates resistance to pyrimethamine and sulfadoxine, respectively. Except for DHPS A437G, the other mentioned mutations are absent in NF54. PAT-039 harbors wild-type (3D7) PfCRT (K76T) and PfMDR1 (N86, Y184, D1246) alleles, identical to NF54 sequences at these loci (Fig. 1d). To perform the genetic cross, mature gametocytes of PAT-039 and NF54 were combined 1:1 and fed to female *Anopheles gambiae* mosquitoes (Fig. 1e). Recombinant sporozoites that subsequently developed in these mosquitoes were transmitted to three human liver–chimeric FRD-NOD mice, either by intravenous inoculation of dissected sporozoites or mosquito bite (Fig. 1e). Progeny pools of ring-stage asexual parasites were obtained from the mice, *in vitro* culture-adapted and used for bulk drug selections, generation of individual progeny clones, and competitive growth assays to identify genetic loci contributing to antimalarial drug susceptibility and fitness (Fig. 1f,g).

**Figure 1.**
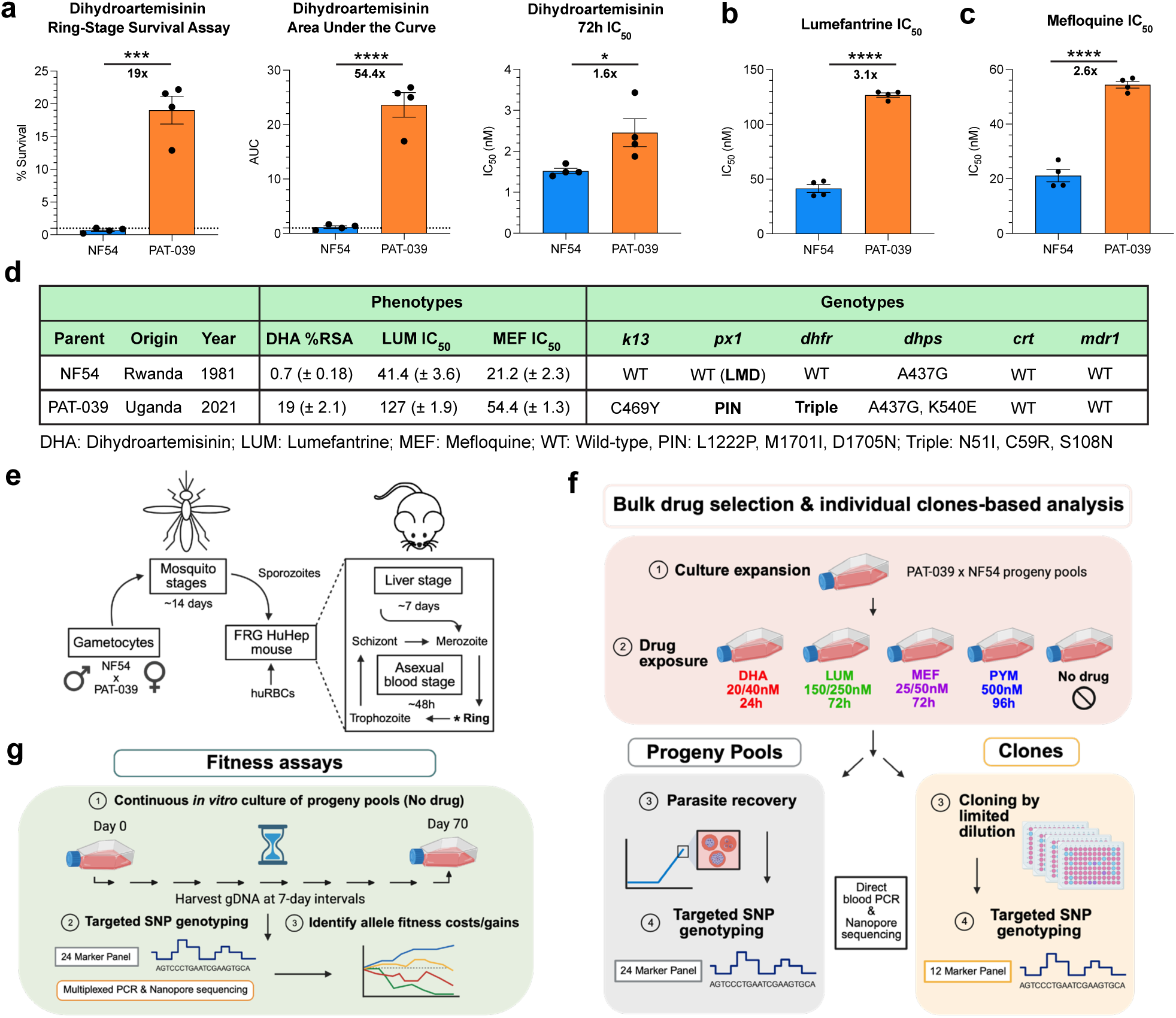
Experimental workflow for the genetic cross of PAT-039 and NF54 parasite lines and genotype-phenotype profiling of progeny using drug selection of progeny pools, clones-based analysis and fitness assays. **a**, Susceptibility of NF54 and PAT-039 cross parents to DHA. Ring-stage survival assays (RSA) were used to calculate percentage survival and area under the curve (AUC). IC_50_ was determined using 72h assays. Susceptibility of cross parents to LUM (**b**) and MEF (**c**) determined using 72h IC_50_ assays. Values represented by bars are the means ± standard error of the mean (SEM) (n = 4). Statistical significance was determined using unpaired t-tests (*P < 0.05, ***P < 0.0005, ****P < 0.00005) and is annotated above bars. The numbers below the bars indicate the IC _50_ fold shifts between parental lines. **d**, Table summarising the origins, drug susceptibility phenotypes and genotypic characteristics across key drug resistance markers of the parental lines used in the genetic cross. DHA RSA values represent percentage survival (%). LUM and MEF IC_50_ values represent nanomolar (nM). **e**, Overview of the experimental genetic cross approach in *Anopheles gambiae* mosquitoes and humanised FRG HuHep mice. huRBCs, human red blood cells. **f**, Pooled drug-selection and clones-based strategy using targeted SNP genotyping of cross progeny to identify genetic markers associated with drug resistance in pools and among individual clones. **g**, Fitness assay approach to identify fitness phenotypes associated with parental allele inheritance. Images were created with BioRender.com.

### PX1 PIN mutations were enriched under dihydroartemisinin or lumefantrine selection

Bulk progeny populations from the PAT-039 × NF54 cross were exposed to intermediate and high concentrations of DHA (20 and 40 nM) and LUM (150 and 250 nM) for 24–72h to select for resistant progeny that could survive under drug pressure (Fig. 1f). Progeny pools were also exposed to MEF (25 and 50 nM), an aminoalcohol structurally related to LUM and thought to share mechanisms of action and/or resistance. The antifolate pyrimethamine (PYM) was included as a positive selection control, as inheritance of the triple-mutant *dhfr* allele from PAT-039 confers high-level PYM resistance and is expected to be strongly enriched in the PYM-selected pool. Following parasite recovery after drug exposure, the drug-selected progeny pools were genotyped using multiplexed PCR and amplicon sequencing targeting 24 loci distributed across the *P. falciparum* genome. These 24 loci encompass 36 single-nucleotide polymorphisms (SNPs) that are present in the PAT-039, but not NF54 strain, and are associated with altered susceptibility to common antimalarials and preclinical candidates, or have housekeeping cellular functions such as DNA replication (Extended Data Fig. 2 displays parental genotypes for each marker in addition to full and abbreviated gene names; additional information is available in Supplementary Fig. 1 and Supplementary Table 2). Targeted genotyping of untreated control samples revealed that the recombinant pools contained a balanced ratio of NF54 and PAT-039 alleles for the large majority of the markers, indicating an even inheritance of alleles among the progeny (Supplementary Fig. 2). Exposure to 500 nM PYM for 96h resulted in complete enrichment of the PAT-039 *dhfr* triple mutant allele, which increased from 50% in the untreated pool to 100% in the PYM-treated sample (Fig. 2a), thus validating the selection and genotyping approach. Selection with DHA, LUM or MEF enriched strongly for the PAT-039 *px1* allele encoding the PIN haplotype. PAT-039 PX1 PIN represented 80-100% of alleles following selection with each of these drugs (Fig 2b,c,d and Supplementary Fig. 2). The relative proportion of PX1 PIN increased by 40%, 53%, and 46% for parasites selected with DHA, LUM and MEF, respectively. In contrast to PX1, enrichment of the mutant C469Y *k13* allele from PAT-039 by selection with DHA was only ∼10% (Fig. 2b and Extended Data Fig. 3). The pronounced enrichment of the PX1 PIN haplotype under selection with all three drugs supports a central role for this locus as a determinant of reduced susceptibility to multiple frontline antimalarials. Multiple other potential markers of altered drug susceptibility were enriched or diminished by various drugs, with similar selection patterns for LUM and MEF (Fig. 2).

**Figure 2.**
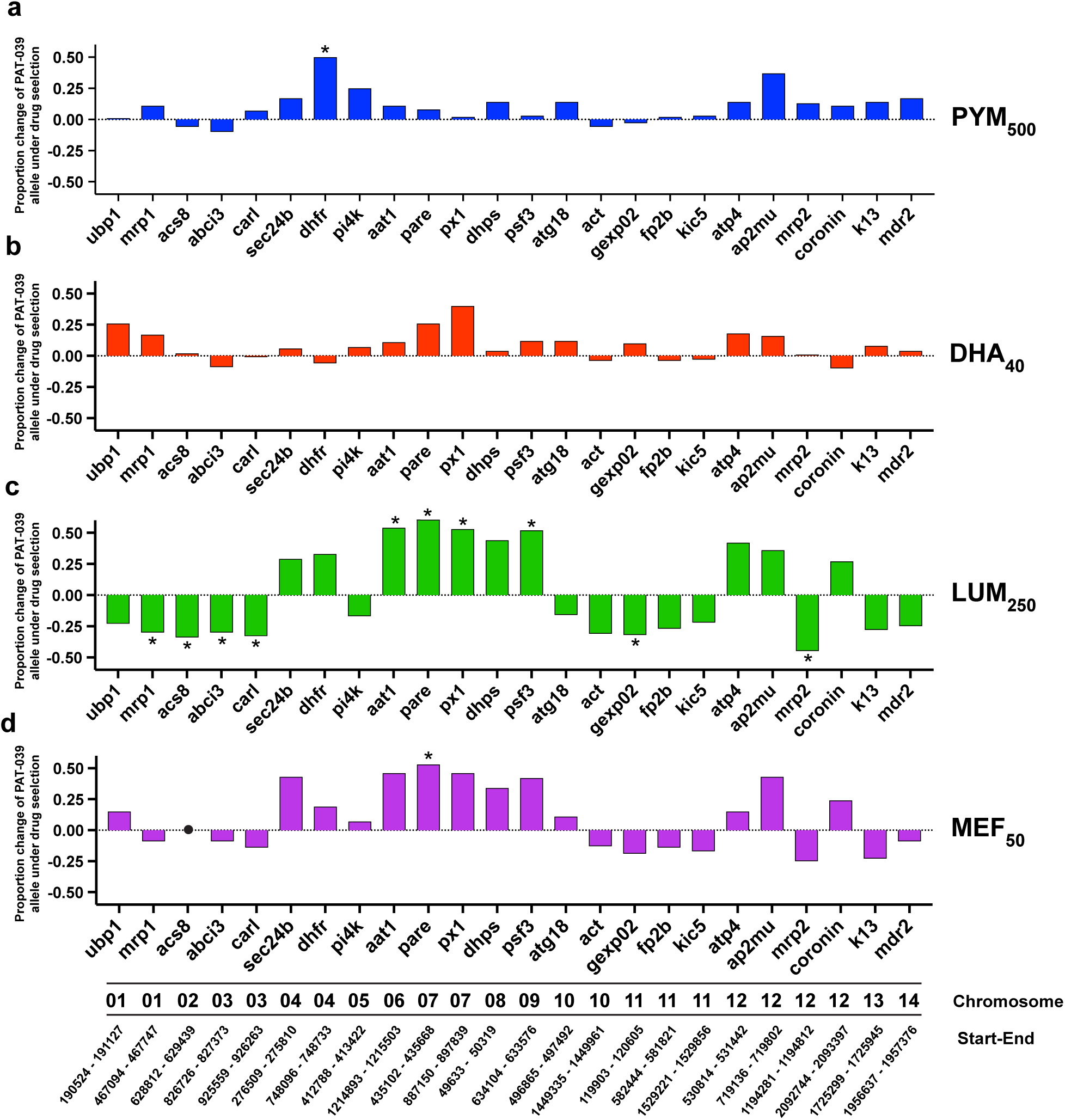
Dihydroartemisinin (DHA) and lumefantrine (LUM) select for the PX1 PIN haplotype in progeny pools. Proportion of the PAT-039 parental allele measured for each gene in the 24-marker SNP genotyping panel following drug selection expressed relative to untreated controls after treatment with 500 nM pyrimethamine (PYM) (**a**), 40 nM DHA (**b**), 250 nM LUM (**c**) and 50 nM MEF (**d**). Each bar represents the proportion of the PAT-039 allele for each marker in drug-selected samples subtracted by the proportion in untreated controls. Bars are colored according to each drug treatment condition. Statistical significance was determined using Z-score (*P < 0.05) and is annotated above bars. Annotated below marker names at the bottom of the figure are the chromosomal locations of each marker and the genomic position of the PCR amplicon used for nanopore sequencing and SNP genotyping. The *acs8* marker was not detected in the MEF_50_ sample and is represented by a black circle. For the full list of marker gene names and IDs, refer to Extended Data Fig. 2.

### Selection with dihydroartemisinin and lumefantrine in progeny converged on distinct genotypic signatures including PX1 PIN

To distinguish clonal genotypes and investigate whether different antimalarial drugs selected for distinct genotypic signatures, we sequenced a subset of 12 marker genes in the progeny clones. Genotyping over 400 individual progeny clones identified 215 genetically unique recombinants. To quantify the effect of drug selection on individual loci, we calculated the proportion of clones carrying the PAT-039 or NF54 allele at each marker for each drug condition relative to untreated controls (Fig. 3a). Across DHA-, LUM-, and MEF-selected clones, the PAT-039 *px1* allele showed the strongest enrichment of any locus examined, increasing in proportion by 40-50% relative to untreated clones (Fig. 3a). All LUM_250_-selected clones harbored the PAT-039 *px1* allele, as did 96% of MEF_50_-selected clones (Supplementary Fig. 3), indicating exceptionally strong selective pressure at this locus. PX1 PIN was also by far the most enriched locus in progeny under selection with DHA (Fig. 3a).

**Figure 3.**
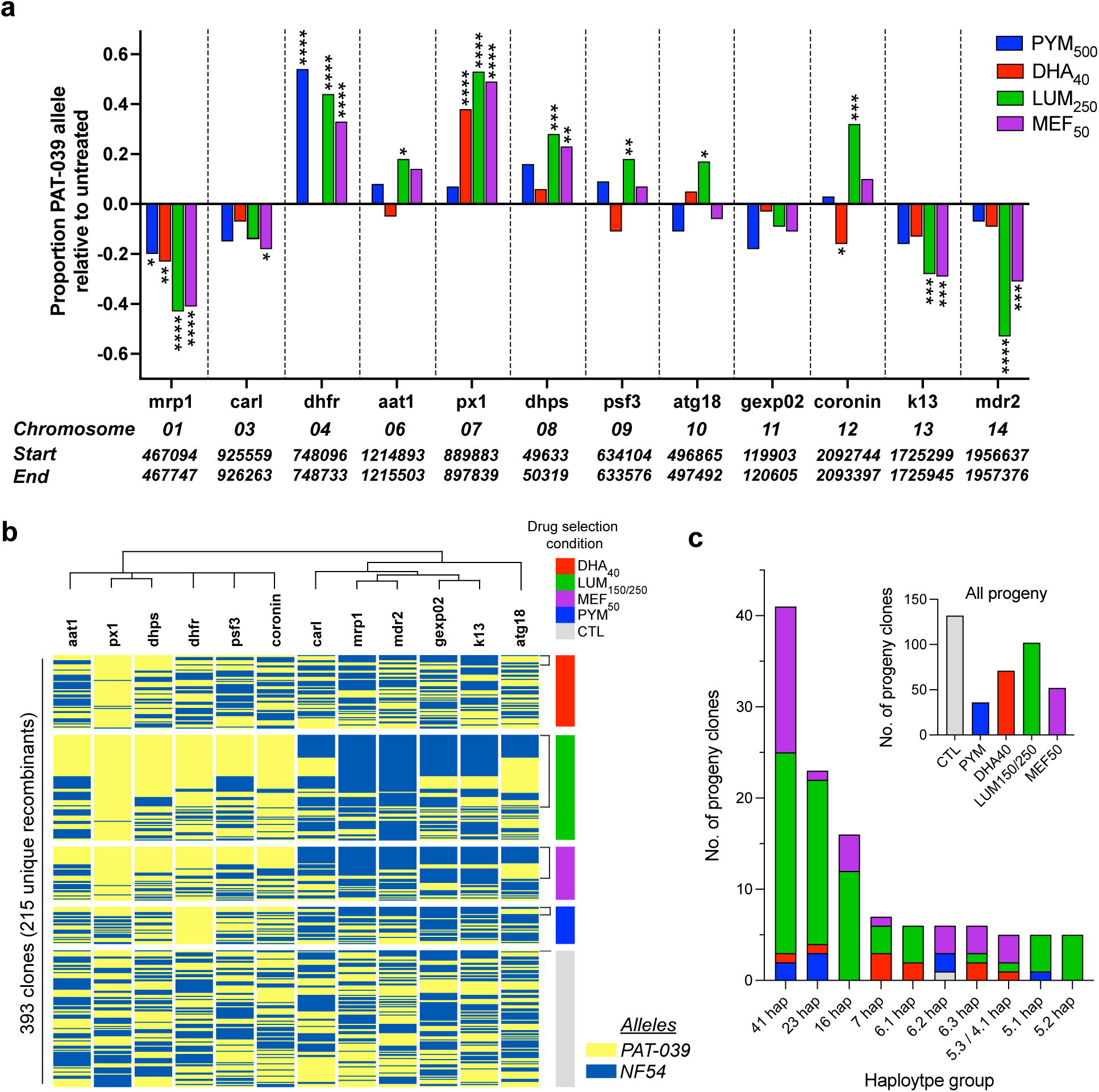
DHA and LUM enrich for clones with specific genotypic signatures including PX1 PIN mutations. **a**, The proportion of clones possessing the PAT-039 parental allele obtained under each drug treatment condition relative to untreated controls for each marker. Bars are colored according to the drug treatment condition from which clones are derived. Annotated below marker names are the chromosomal locations of each marker and the genomic position of the PCR amplicon used for nanopore sequencing and SNP genotyping. Statistical significance was determined using Fisher’s Exact Test on raw counts (*P < 0.05, **P < 0.005, ***P < 0.0005, ****P < 0.00005) and is annotated above bars. **b**, Allelic map of progeny clones showing the parental allele inheritance among the 12 genes in the SNP marker panel across all 393 progeny clones. Progeny are ordered by drug-selection condition from which they were derived, as shown by colored bars. Alleles are colored according to the parental origin (NF54 or PAT-039). Brackets annotate progeny which belong to the 10 largest haplotypic groups. **c**, The number of progeny clones from each drug-selection condition or control comprising the 10 largest haplotypic groups. Bars are colored according to the drug-selection treatment condition from which the clones were derived. The inset graph displays the total number of progeny clones generated under each selection condition. Haplotypic groups are named according to the number of clones constituting each group, where letters are used to distinguish groups with equal numbers of clones. For the full list of marker gene names and IDs, refer to Extended Data Fig. 2. LUM and MEF selections led to enrichment of antifolate resistance-linked *dhfr* and *dhps* alleles in up to 90% of clones and co-inherited with PX1 PIN.

To determine how locus-level selection translated into clonal genotypes, we examined the allelic composition of each progeny clone under each drug selection condition. While we identified several dominant haplotypic groups that emerged under selection with DHA, LUM or PYM, the vast majority of clonal genotypes were derived from a single drug exposure condition (Fig. 3b,c and Extended Data Fig. 4). In spite of the clonal genotypic diversity between selection conditions, nearly all progeny recovered following exposure to all three drugs harbored the PAT-039 *px1* allele (Fig. 3b). In contrast, the PAT-039 C469Y K13 allele was not enriched in clones obtained following DHA selection; instead, a moderate (∼15%) decrease in the relative proportion of the mutant allele was observed after DHA selection (Fig. 3a,b). These findings indicate that, despite drug-specific effects at other loci, DHA, LUM and MEF robustly converge on PX1 PIN as the dominant shared genetic determinant associated with reduced susceptibility to all three antimalarials.

### The PX1 PIN allele conferred a fitness advantage among recombinant progeny

Having identified PX1 as the top marker associated with decreased susceptibility to DHA and LUM, we next assessed the impact of PX1 PIN mutations on parasite asexual fitness using competitive growth assays. To track temporal changes in parental allele frequencies and infer relative fitness effects, bulk progeny pools containing more than 200 recombinants were cultured in the absence of drug pressure for 70 days and genotyped at regular intervals using a 24-marker panel (Fig. 1g, Fig. 4 and Extended Data Fig. 5). We observed a striking increase in the proportion of the PAT-039 *px1* allele, which reached ∼90% of the population by day 63, exceeding that of all other markers (Fig. 4a). We next quantified locus-specific fitness effects by modelling changes in allele frequencies over time to estimate relative fitness coefficients for each marker (Fig. 4b and Supplementary Table 3). Consistent with the growth trends, the PAT-039 *px1* allele was the only locus to show a statistically significant fitness gain (selection coefficient = 0.036), unlike other loci which associated with neutral (*atp4*, *k13*) *or* negative (*ubp1*, *dhfr*, *pi4k*) fitness effects (Fig. 4a,b). This indicates a strong intrinsic fitness advantage conferred by the PX1 PIN haplotype in the absence of drug pressure.

**Figure 4.**
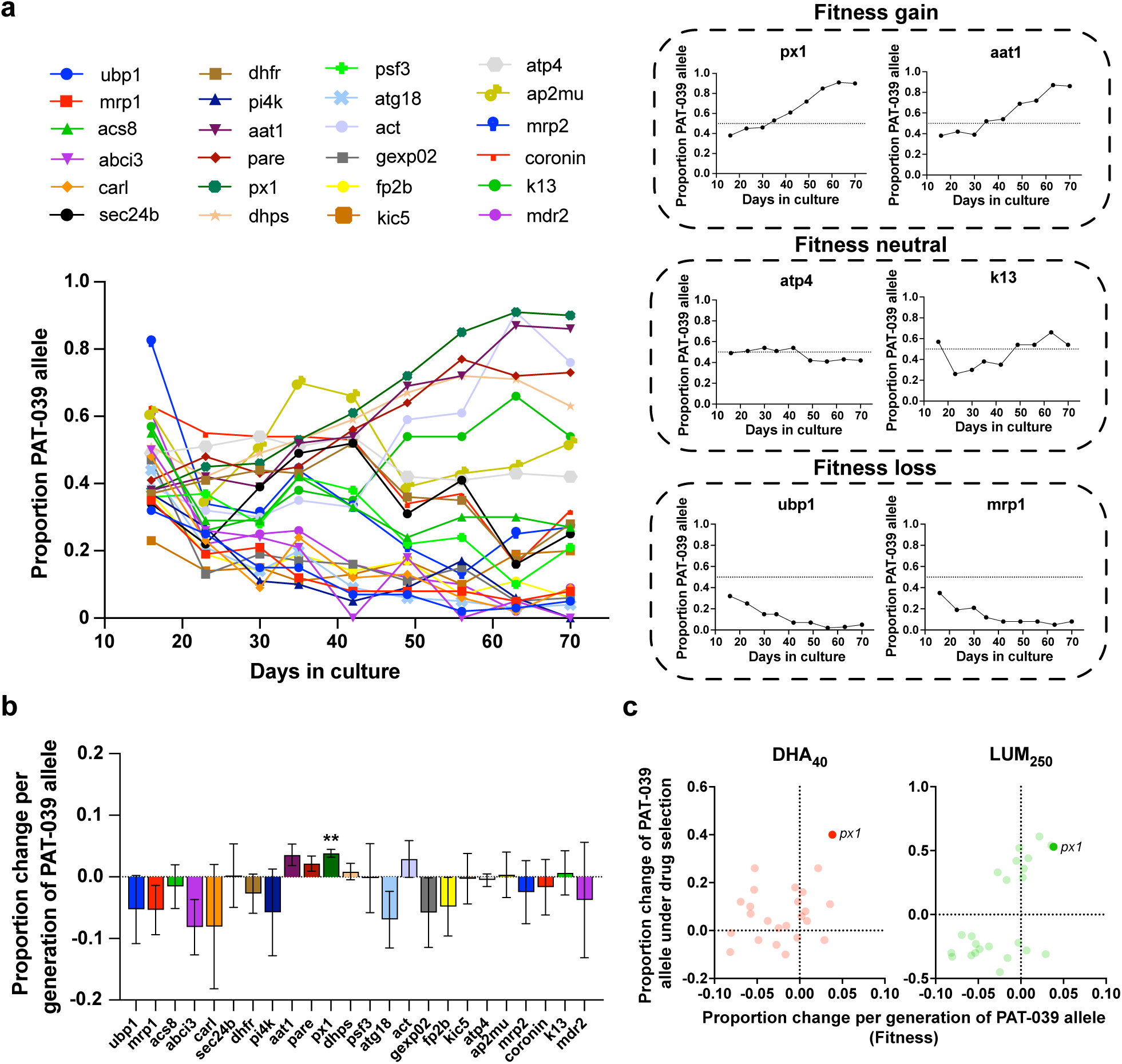
The PX1 PIN allele confers a fitness advantage among recombinant progeny. **a**, Proportion of the PAT-039 allele in progeny pools for all markers in the 24-marker panel measured at nine timepoints across 70 days in culture, in the absence of drug pressure. Points and lines are colored according to each marker. Graphs on the right panel show exemplar markers for which the PAT-039 allele confers either a fitness gain, neutrality or loss. **b**, Percentage change per generation of the PAT-039 allele for each marker determined between each timepoint in the fitness assay and averaged across all timepoints. Values represent the means ± SEM (n = 8). Statistical significance was determined using one sample t-tests (**P < 0.005) and is annotated above bars. Marker genes are arranged in ascending chromosomal order. For the full list of marker gene names and IDs, refer to Extended Data Fig. 2. **c**, Association between the fitness phenotype (data shown in Fig. 4b) and drug-selection phenotype (data shown in Fig. 2) under DHA and LUM selection. Points represent individual marker genes colored according to drug selection condition. Data points for *px1* are annotated.

We compared locus-specific fitness effects with changes in PAT-039 allele frequency for each drug. PX1 PIN displayed exceptional fitness in both the presence and absence of DHA and LUM, unlike all other polymorphisms studied herein (Fig. 4c and Supplementary Fig. 4). In contrast, drug resistance-associated transporters including *abci3, mrp1, mrp2* and *mdr2* in which the mutant alleles were selected against by LUM pressure, exhibited fitness defects (Fig. 4b,c and Extended Data Fig. 5).

### Dihydroartemisinin and lumefantrine selection enriched for the PX1 PIN haplotype in Ugandan clinical isolates

To determine whether the enrichment of PX1 mutations under DHA, LUM and MEF pressure were also seen in other parasite genetic backgrounds, we applied a bulk drug-selection approach to a pool of 18 distinct *P. falciparum* isolates collected from northern and eastern Uganda (Fig. 5). We exposed this pool to DHA for 24h, to LUM or MEF for 72h, or to no drug (Fig. 5a,b and Supplementary Table 4). To identify SNPs enriched by drug selection, allele frequencies of the 24-marker panel were quantified and compared (Extended Data Fig. 6 and Supplementary Fig. 5).

**Figure 5.**
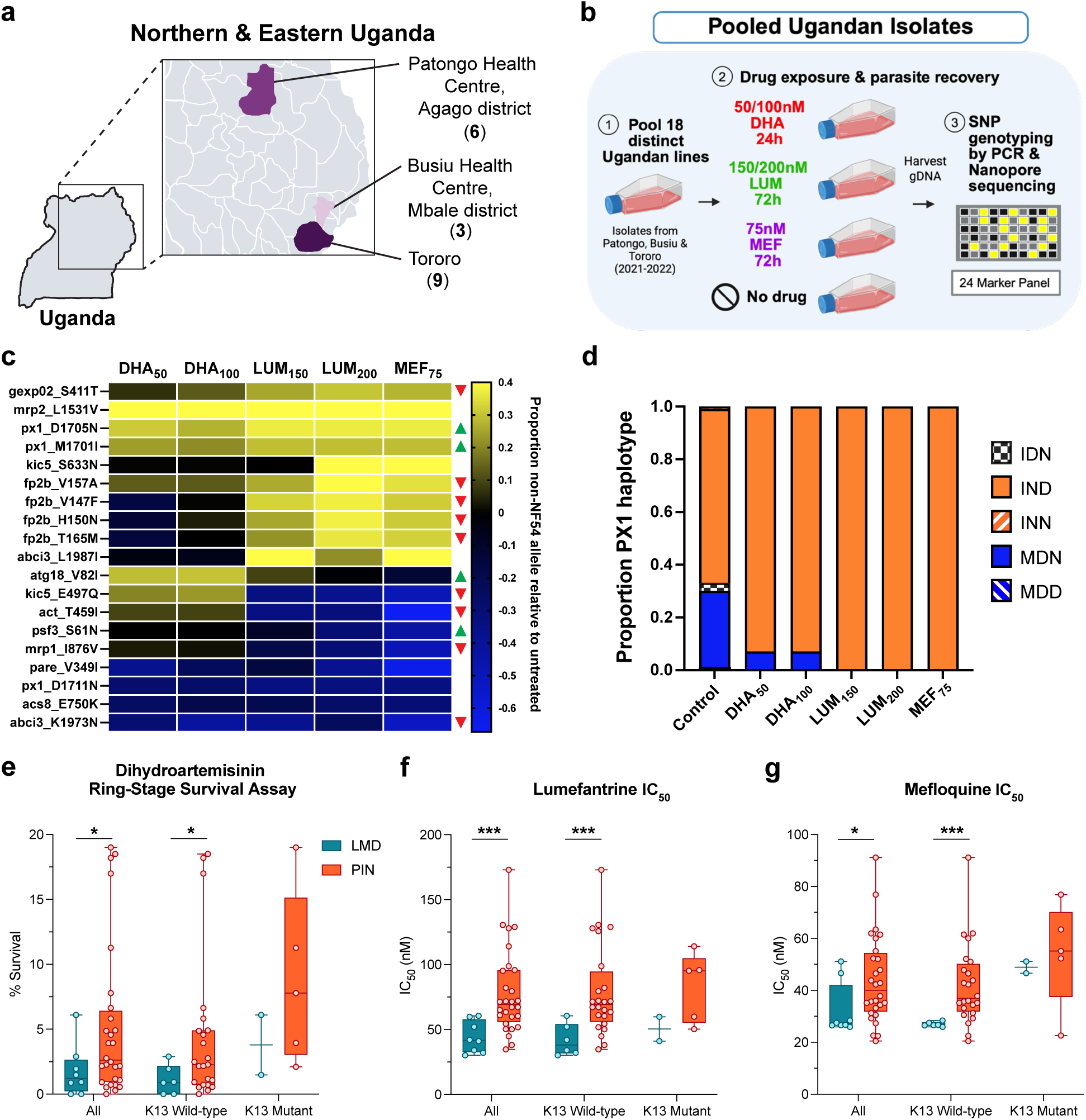
Selection of a pool of 18 Ugandan clinical isolates with DHA and LUM enriches for PX1 PIN mutations. **a**, Map of northern and eastern Uganda colored and annotated by district from which clinical isolates used in pooled drug-selection assays were collected. Numbers in brackets represent the number of isolates derived from each district. **b**, Experimental approach for drug selection and SNP marker genotyping of pooled Ugandan clinical isolates. Image was created with BioRender.com. **c**, Heat map displaying the change in the proportion of non-NF54 alleles measured in pooled Ugandan isolates under different drug-selection conditions relative to untreated controls. The yellow-blue scale bar represents an increase or decrease in the proportion of non-NF54 allele in the drug-treated samples compared to the untreated control. Colored triangles denote whether the proportion of the allele that is also present in PAT-039 displays an increase (green) or decrease (red) in drug-selected cross progeny (refer to Fig. 2). For the full list of marker gene names and IDs, refer to Extended Data Fig. 2. **d**, Proportion of each PX1 haplotype in Ugandan isolate pools under control (untreated) or drug-treated conditions. Haplotypes are classified according to the presence of the wild-type NF54 (MDD) or mutant alleles (IDN, IND, INN, MDN) at the M1701I, D1705N and D1711N positions of PX1. Bars are colored according to each haplotype. **e-g,** Drug susceptibility phenotypes determined for a panel of 32 Ugandan clinical isolates grouped according to PX1 genotype (LMD or PIN) and K13 genotype (wild-type or mutant). K13 mutant group harbors C469Y or A675 mutations whereas WT group harbors either full-length wild-type or non-WHO-validated K13 mutations. **e.** Percentage ring-stage survival determined following exposure to DHA. **f**, LUM IC_50_ and (**g**) MEF IC_50_. Lines in all graphs represent experimental means across all Ugandan isolates profiled in the assays. Error bars represent interquartile range. Individual points represent the mean value determined across at least three biological replicates for each individual isolate. Statistical significance was determined using unpaired t-tests (*P < 0.05, ***P < 0.0005, ****P < 0.00005) and is annotated above bars.

The PX1 M1701I and D1705N SNPs and one other polymorphism (the *mrp2* L1531V mutation) were the only SNPs enriched consistently, and these ranked among the most strongly enriched variants in all drug treatment conditions (reaching 90-100% prevalence), including multiple concentrations of both DHA and LUM (Fig. 5c, Extended Data Fig. 6 and Supplementary Fig. 5). Sequencing of the full length *px1* gene revealed enrichment of the L1222P mutation, which indicates near fixation of the PIN haplotype under drug selection (Extended Data Fig. 7). An additional PX1 mutation present in the Ugandan clinical isolates pool, but absent from the PAT-039 cross parent, D1711N, was selected against under exposure to DHA, LUM and MEF, consistent with a specific selection for the PIN haplotype rather than PX1 polymorphisms more broadly (Fig. 5d). These data demonstrate that PX1 PIN is robustly selected by DHA and LUM in African parasites.

### The PX1 PIN haplotype associates with reduced *in vitro* susceptibility to dihydroartemisinin and lumefantrine among Ugandan clinical isolates

Since PX1 PIN has been shown to be the top candidate marker for reduced DHA and LUM susceptibility, we next assessed the relationship between the PX1 PIN haplotype (L1222P, M1701I, D1705N) and drug susceptibility in a panel of 32 culture-adapted genetically diverse Ugandan clinical isolates. In DHA ring-stage survival assays, PIN isolates exhibited significantly higher mean survival compared with wild-type isolates (Fig. 5e). In growth inhibition assays for LUM and MEF, PIN isolates displayed significantly elevated IC₅₀ values (Fig. 5f,g and Supplementary Table 5). These results establish a strong association between the PX1 PIN haplotype and reduced susceptibility to DHA, LUM and MEF in Ugandan parasites.

Substantial heterogeneity in drug susceptibility was observed among PIN isolates, suggesting that additional genetic factors may modulate drug susceptibility. To explore potential interactions with known ART-R determinants, we further stratified drug responses by *k13* genotype (Fig. 5e-g, Supplementary Table 5). The absence of PX1 and K13 mutations invariably conferred parasite sensitivity to all three antimalarials (Fig. 5e-g). Isolates carrying both PX1 PIN and K13 mutations exhibited higher DHA RSA survival, as well as increased LUM and MEF IC₅₀ values, indicating potential additive effects between *px1* and *k13* polymorphisms on reduced drug sensitivity. Nonetheless, isolates with the PX1 PIN haplotype without K13 mutations displayed reduced susceptibility to DHA, LUM and MEF, suggesting that PX1 mutations can confer increased tolerance to these antimalarials independently of K13.

## Discussion

Leveraging the first experimental genetic cross including a recent Ugandan *P. falciparum* isolate, we dissected the genetic basis of reduced susceptibility to Africa’s frontline ACT, artemether–lumefantrine. Applying a targeted genotyping approach to drug-exposed bulk progeny pools and progeny clones, and performing competitive fitness assays, we showed that the phosphoinositide-binding protein PX1 PIN haplotype was associated with reduced sensitivity to DHA and LUM. This association was independently validated in genetically diverse Ugandan clinical isolates, with *in vitro* drug assays demonstrating that PIN-carrying parasites exhibited reduced susceptibility to DHA, LUM, and MEF and that selection with these drugs enriched the PIN haplotype. These results establish PX1 PIN as a novel marker of reduced *in vitro* DHA and LUM sensitivity and thus as an emerging multidrug resistance locus. Our findings from competitive growth assays also indicated that the PX1 PIN haplotype confers enhanced parasite fitness. Overall, our findings from a genetic cross and a recent population-based report from Uganda^32^, indicate that the PX1 PIN haplotype which has rapidly expanded in Uganda, is associated with reduced susceptibility to both components of AL.

Beyond establishing PX1 PIN as a resistance marker, our findings raise questions regarding its physiological function and mechanistic role in antimalarial drug resistance. The *px1* gene encodes a phosphoinositide-binding protein that localizes to the parasite digestive vacuole membrane and has been implicated in hemoglobin trafficking^39^. K13 propeller domain mutations are believed to mediate ART-R via disruption of hemoglobin trafficking. Mutations in PX1 may similarly perturb hemoglobin uptake or processing, reducing hemoglobin turnover and free heme availability, paralleling proposed mechanisms of K13-mediated ART-R^8^. Such changes could also influence susceptibility to ACT partner drugs, including lumefantrine, mefloquine, piperaquine and amodiaquine. LUM acts either by disrupting heme detoxification within the digestive vacuole or by targeting an as-yet unidentified cytosolic factor^40,41^. The PX1 PIN mutations may reduce LUM sensitivity by altering vacuolar physiology and the levels of hemoglobin or its processing products or may reduce access of LUM to its cytosolic target. Dissecting the mechanism underpinning *px1* polymorphisms in resistance to distinct chemical classes will require targeted genetic manipulation, biochemical characterization and cell biological analyses of PX1 function. Further evaluation of PX1 variants in the absence and presence of K13 mutations will help to determine whether these loci act independently or convergently to modulate drug susceptibility.

Unexpectedly, DHA selection applied to cross progeny and Ugandan isolates did not enrich for K13 mutations, despite associations between variants such as C469Y and A675V with reduced DHA susceptibility in Uganda^17,24^. Our findings suggest K13 may not be the primary artemisinin resistance determinant in the East African genetic background. In Southeast Asian parasites, K13 mutations confer early ring-stage tolerance without altering IC_50_ values to DHA in asexual blood stage growth inhibition assays^42^. This paradigm may not translate to African parasites, implying additional resistance mechanisms^11^. The absence of K13 enrichment by DHA may reflect the 24h DHA drug exposure regimen employed in this study. While K13*-*mediated resistance is most evident following short pulses that mimic *in vivo* exposure, a longer duration of drug exposure may select for distinct DHA resistance mechanisms. While artemisinin derivatives are rapidly cleared from the plasma, evidence suggests a longer elimination half-life in red blood cells^43^. This prolonged intracellular effect and the fact that AL is administered twice-daily^44^, versus once daily for all other ACTs, may subject the parasite to DHA pressure over much of the intraerythrocytic cycle and render resistance mechanisms limited to early ring stages less beneficial for parasite survival. Systematic studies across parasite developmental stages and drug exposure durations will be critical to better characterize resistance mechanisms.

LUM and MEF exposure exerted a strong selective effect against PAT-039 mutant alleles for genes encoding the putative transporters *abci3, mrp1, mrp2* and *mdr2* in progeny clones and bulk progeny pools, indicating that wild-type transporter function is favoured under exposure to these drugs, These mutations present in the PAT-039 parent are also linked to growth defects in our fitness assays, suggesting that these loci may lead to negative effects on parasite physiology, possibly affecting drug uptake or vacuolar homeostasis^40,45^. The predominance of wild-type alleles in *mrp1* and other transporter proteins such as *pfcrt* and *pfmdr1* in East African parasite populations supports this interpretation^46,47^. Surprisingly, certain haplotypic groups emerged under selection with DHA, LUM or PYM indicating that diverse antimalarials can enrich for the same haplotypes. The co-inheritance of mutant *dhfr* alleles under LUM and MEF selection raises the possibility that parasites carrying PX1 mutations may simultaneously tolerate antifolate pressure through the acquisition of a shared genotype, suggesting that selection for resistance to one drug class may indirectly stabilize resistance alleles to others.

While our targeted amplicon sequencing strategy enabled sensitive and scalable detection of candidate resistance loci across progeny pools, it prioritized known polymorphic genes with established links to antimalarial resistance. Whole-genome analyses of bulk progeny and clones may identify additional drug susceptibility modifiers and/or epistatic interactions. Consistent with this, substantial phenotypic variability among PIN-carrying clinical isolates indicates that PX1 polymorphisms operate within a broader polygenic landscape. Defining this architecture will be critical for anticipating resistance trajectories and informing optimal antimalarial treatment policies.

In this study, we combined orthogonal approaches including an experimental cross and the drug selection of pooled clinical isolates to converge on *px1* polymorphisms as a mechanism of decreased susceptibility to multiple first line antimalarials. These findings argue strongly for the incorporation of *px1* into existing genomic surveillance efforts and raise the possibility that monitoring *px1* dynamics could provide early warning of declining AL efficacy.

## Methods

### In vitro *P. falciparum* cell culture

*P. falciparum* asexual blood stage parasites were cultured in human A+ RBCs obtained from the New York Blood Center (New York, NY) at 3% hematocrit using RPMI 1640 medium supplemented with 25 mM Hepes, sodium bicarbonate (2.1 g/liter), gentamicin (10 μg/ml), 50 μM hypoxanthine, 0.5% (w/v) AlbuMAX II (Gibco), and 10% (v/v) human serum (Grifols). Parasites were maintained at 37°C under 5% O2/5% CO2/90% N2 gas conditions.

### Parasite lines

*P. falciparum* isolates were collected from human patient samples in Uganda and thereafter subjected to short-term *in vitro* culture adaptation. Routine testing and removal of mycoplasma by sparfloxacin treatment was performed as described to ensure cultures were mycoplasma-free if any were found to be mycoplasma-positive^1^.

### Genetic cross

The *P. falciparum* PAT-039 isolate was collected at Patongo Health Centre III, Agago District, Uganda in 2021 and the NF54 isolate was provided by Johns Hopkins University^2^. As there is often a high multiplicity of infection in African lines, we derived a clone of the PAT-039 parent by limiting dilution which is described in the section below. The clone of PAT-039 was used in the experimental cross. PAT-039 and NF54 gametocytes were cultured as described previously^3,4^. Briefly, mature gametocyte cultures (days 15–18) were normalized to 0.6% gametocytemia, mixed 1:1, and fed to *Anopheles gambiae* mosquitoes. Oocyst prevalence and intensity were assessed by mercurochrome staining and brightfield microscopy of midguts using a Nikon E600 microscope and PlanApo 10× objective. Salivary gland sporozoites were quantified by hemocytometer counts from homogenizing glands of 20 mosquitoes.

Animal experiments were approved by the Johns Hopkins IACUC (protocol MO23H231). HuHep FRG-NOD mice with ≥70% human hepatocyte repopulation were purchased from Yecuris and housed with daily monitoring. Two independent mosquito infection cycles were performed. In cycle 1, oocyst prevalence was 50% (mean 7 oocysts/mosquito), with a mean of 21,333 salivary gland sporozoites per mosquito. A total of 500,000 sporozoites were intravenously injected into two mice each. In cycle 2, oocyst prevalence was 94% (mean 22 oocysts/mosquito), with a mean of 18,919 sporozoites per mosquito. One mouse was infected via 50–60 mosquito bites and another by intravenous injection of 1,000,000 sporozoites. Mice infected by intravenous sporozoite injection received prophylactic intraperitoneal injections of 1,000 U penicillin and 1 mg streptomycin for four days. To support the transition from liver to blood stage infection, 500 µL of 70% human RBCs were administered intraperitoneally on days 5, 6 and 7 post-infection. Parasitemia was assessed at day 7.5 and ranged between 0.05%-0.7%.

### Drug selections in bulk progeny pools

Bulk progeny pools were *in vitro* culture adapted and subjected to drug exposures at 3% hematocrit and 1% parasitemia in 10 mL cultures as follows: 20 or 40 nM DHA for 24h (10-20× IC_50_ for NF54), 150 or 250 nM LUM for 72h (4-7× IC_50_ for NF54), 25 or 50 nM MEF for 72h (1.5-3× IC_50_ for NF54), and 500 nM PYM for 96h (27× IC_50_ for NF54). Drug-medium was replenished every 24h and subsequently removed following 24, 72 or 96h of exposure by centrifuging the culture at 1500 rpm for 5 min, discarding the supernatant and washing with 50 ml of fresh media. Cultures were incubated at 37°C and monitored daily. Once parasitemia reached 1%, which varied between 2 to 16 days from the start of treatment, gDNA was harvested from the cultures.

### Clinical isolates and progeny cloning

The Ugandan isolate clones and cross progeny clones were isolated respectively from the bulk parent cultures or bulk progeny pools by limiting dilution at a density of 0.8 parasites per well. Cultures were maintained with medium changes twice weekly, and fresh red blood cells were added once per week. From day 16, wells were screened for parasitemia growth by flow cytometry and positive wells were expanded for SNP genotyping by PCR and nanopore sequencing as described below.

### Fitness assays

The bulk progeny pool was subjected to *in vitro* culture adaptation and maintained at 3% hematocrit and 2-4% parasitemia for up to 70 days post mouse sacrifice. Parasites were harvested once every week for gDNA in order to perform SNP genotyping as described below.

### *In vitro* drug response assays

#### Dihydroartemisinin, lumefantrine and mefloquine

To obtain tightly synchronized parasites for determining RSA and DHA IC_50_ values, cultures were treated with 5% D-Sorbitol (Sigma-Aldrich) for 15 min at 37**°**C to remove mature parasites, we then cultured for 40h and purified late-stage segmented schizonts over 35 and 65% Percoll double density gradients^5^. Purified schizonts were incubated with fresh RBCs for 3h and treated with 5% w/v D-Sorbitol to obtain 0- to 3h post-invasion early rings. These rings were exposed to the DHA concentration of 700 nM for RSA and 10-point serially diluted two-fold concentrations starting from 100 nM for DHA 16h and 72h IC_50_ assays in 96-well plates. Wells were inoculated with 200-μl samples (at 0.9% starting parasitemia and 1% hematocrit), with technical duplicates. Untreated control wells were included alongside. For RSAs and 16h IC_50_ assays, after a 4h and 16h incubation at 37**°**C, wells were washed three times with complete medium to remove drug, and contents were transferred to fresh 96-well plates using the Freedom EVO MCA96 liquid-handling instrument (Tecan). Cultures were subsequently maintained for an additional 68h and 56h in drug-free medium, respectively.

Standardised 72h dose-response assays were conducted for LUM and MEF. Sorbitol-synchronised ring-stage parasites were exposed for 72h at 10-point serially diluted concentrations starting from 1200 nM and 400 nM for LUM and MEF, respectively. Untreated control wells were run in parallel. Dose response assays were performed in at least three independent experiments with technical duplicates.

### Flow cytometry for quantification of viable parasites

Parasitemia for drug-treated and untreated wells were measured at 72h by flow cytometry, as previously described^6^. Briefly, parasites were incubated with 1x SYBR Green (ThermoFisher Scientific) and 100 nM MitoTracker DeepRed (Thermo Fisher Scientific) for 40 min at 37°C, followed by quenching with 1x PBS. On average, we analyzed 10,000 cells per sample, using an iQue Screener Plus cytometer (Sartorius). Viable parasites were defined as the percentage of MitoTracker-positive and SYBR Green-positive cells.

### Calculation of RSA and IC_50_ values

For all drug assays conducted here, we included kill controls in which 1.5 μM DHA-treated parasites were used as a background control to achieve complete parasite killing and subtracted this percent parasitemia from the parasitemia measured for each well. Parasite survival in the presence of DHA was expressed as the percentage of the background-subtracted parasitemia in the 700 nM DHA-treated samples divided by the background-subtracted parasitemia of the untreated samples. Mean RSA survival rates of >1% were defined as DHA resistant^7^. IC_50_ values were determined by applying a nonlinear regression model (sigmoidal dose-response with variable slope) on the normalized percent survival across the log-transformed drug concentrations, using Prism v10.4.1 (GraphPad).

### Drug selections in Ugandan isolate pools

The 18 Ugandan parasite isolates were normalized to 1% parasitemia and mixed in equal ratios to establish the pool named as “UGpool”. The parasite pool was then divided into 5mL, 1% parasitemia, 2% hematocrit cultures and exposed to 50 nM or 100 nM DHA, 150 nM or 200 nM LUM, or 75 nM MEF for 72h or in the absence of drug. Following drug removal by washing with 50 mL of no-drug media and centrifuging at 1500 rpm for 5 min, parasites were then placed back into culture and allowed to recover. Once the drug-treated UGpools reached 1% parasitemia, samples were harvested for genomic DNA. Drug-treated UGpools were harvested 16-19 days after the start of DHA exposure, 22-23 days post-LUM treatment, and 21 days post-MEF treatment. The untreated UGpool was also harvested for gDNA at the initiation of the drug-treatments, maintained alongside and subsequently harvested at a similar time point as when drug-treated samples were collected. Genomic DNA extraction was performed using the Qiagen Blood Mini kit according to the manufacturer’s instructions. DHA-treated and recovered parasites from the UGpool were also subsequently cloned out by limiting dilution for further drug phenotyping experiments and genotyped for *mdr1* and *k13* by PCR and Sanger sequencing as described below.

### Progeny genotyping

#### SNP marker panel design

Genes represented in the 24-marker genotyping panel were selected on the basis of possessing SNPs in the parental PAT-039 genome that were absent in NF54, and represented either validated drug resistance markers or housekeeping genes sampled from across all 14 *P. falciparum* chromosomes. Primers to amplify marker genes by PCR were designed to adhere to the following criteria; amplicon size of ∼600bp encompassing at least one SNP, primer GC content of 40-60% and melting temperature of 55-58°C. Primer sequences were analysed using ThermoFisher Scientific Multiple Primer Analyzer to identify potential cross-dimer interactions prior to pooling for multiplexed PCR. Primers were arranged into the following seven pools with minimal cross-binding; Pool 1 (*k13*, *dhfr*, *px1*), Pool 2 (*atg18*, *dhps*, *gexp02*), Pool 3 (*mrp1*, *coronin*, *aat1*), Pool 4 (*carl*, *mdr2*, *psf3*), Pool 5 (*pare*, *fp2b*, *ubp1*, *kic5*), Pool 6 (*abci3*, *acs8*, *sec24b*), Pool 7 (*act*, *pi4k*, *mrp2*), Pool 8 (*ap2mu*, *atp4*).

### Multiplex PCR

All PCR reactions were performed in 15 µL final reaction volumes including the following components; 1 µL infected red blood cells (iRBC) template (minimum parasitemia 0.2%), 1x CloneAmp Hifi PCR Premix (Takara), 0.4-0.6 µM premixed primer pair and nuclease-free water up to 15 µL. PCR reactions were mixed by agitation and centrifugation to resuspend iRBCs completely prior to thermocycling using the following conditions; 95 °C for 3 min, followed by 35 cycles of 98 °C for 10 sec, 55 °C for 15 sec and 62 °C for 15 sec. PCR reactions were pelleted by centrifugation and 10 uL of supernatant was pooled from each multiplexed PCR reaction and purified using the Monarch® PCR & DNA Cleanup Kit (NEB) with a final elution volume of 15 uL. Purified DNA was quantified by NanoDrop and diluted to 600 ng/uL prior to submission to Plasmidsaurus Premium PCR service for Oxford Nanopore sequencing.

### Nanopore sequence analysis and variant calling

Following pooled amplicon sequencing, fastq files were processed using the nano-rave (Nanopore Rapid Analysis and Variant Explorer) analysis pipeline via Nextflow^8^. Following QC metrics, sequence reads were mapped against 3D7 reference sequences for each of the amplicon target sequences using minimap2, after which .sam files were converted to .bam files using samtools. ‘Freebayes’ was used for variant calling to generate VCF outputs and to calculate SNP allele frequencies. IGV was used to verify the allele frequencies for each SNP in the target amplicon regions.

### Data Availability

Sequence data that support the findings of this study have been deposited in NCBI SRA under BioProject PRJNA1390316. All other data supporting the findings of this study are available within the paper and its supplementary information files or can be requested for access.

## Acknowledgements

This work was supported by National Institutes of Health (NIH) National Institute of Allergy and Infectious Diseases (NIAID) grants R01AI182318 to SM, AI075045 to PJR, and AI173557 to MDC. We thank David Fidock for his support and acknowledge the Center for Malaria Therapeutics and Antimicrobial Resistance and the Aaron Diamond AIDS Research Center, Division of Infectious Diseases, Department of Medicine, Columbia University Vagelos College of Physicians and Surgeons, for providing equipment and resources. We also acknowledge the support of the Division of Infectious Diseases at Columbia University. We are grateful to Bloomberg Philanthropies for supporting the insectary and parasitology core facilities at the Johns Hopkins Malaria Research Institute. We thank Jeff Bailey and Jonathan Juliano for helpful discussions.

## Author Contributions

CABL and AJ performed research and data analyses and provided intellectual contributions to the study. SK assessed the mosquito infections and performed the experiments in the HuHep mice. AT and GM generated the gametocyte cultures and performed the mosquito feeds. AAMG and LR contributed to PCR Sanger sequencing and *in vitro* culture adaptation. SO and MO collected and culture-adapted field samples. KN contributed to *px1* genotyping methods. PR and MC supervised the collection of Ugandan field samples. PS supervised research on mosquito and mouse work for the genetic crosses. SM designed the study and supervised the overall research project. CABL, AJ and SM wrote the manuscript, with input from all authors.

## Competing interests

The authors declare no competing interests.

## Materials & Correspondence

Correspondence to Sachel Mok.

## Additional Information

Extended data can be found in the ‘Extended Data Figures’ PDF file. Supplementary figures and tables can be found in the ‘Supplementary Figures & Tables’ PDF file.

## Notes

### Competing Interest Statement

The authors have declared no competing interest.

### Summary of Updates

This version has the Acknowledgements section updated.

