## Extended Data Figures for "A *Plasmodium falciparum* PX1 haplotype is associated with reduced susceptibility to artemisinin and lumefantrine"

This file contains Extended Data Figures 1-7

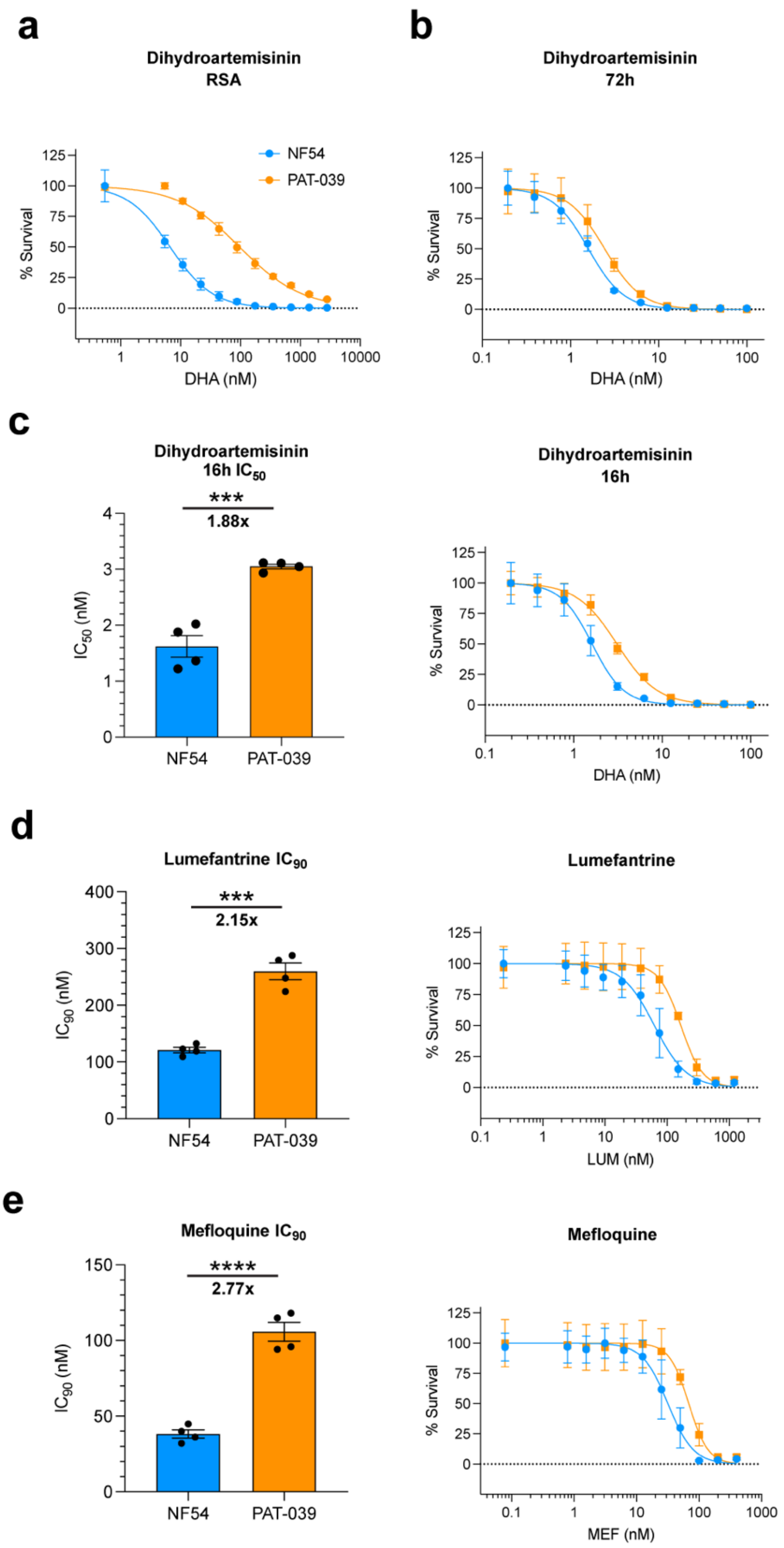


**Extended Data Figure 1. Full drug susceptibility phenotypes of the cross parents PAT-039 and NF54 to dihydroartemisinin (DHA), lumefantrine (LUM) and mefloquine (MEF). a**, Dose-response curves for cross parent lines exposed to serial dilutions of DHA in ring-stage survival assays. **b**, Dose-response curves determined from 72h exposure to DHA in $IC_{50}$ assays. **c**, $IC_{50}$ values and dose-response curves determined from 16h exposure to DHA. **d**, $IC_{90}$ values and dose-response curves determined from 72h exposure to LUM. **e**, $IC_{90}$ values and dose-response curves determined from 72h exposure to MEF. Each line in dose-response curve graphs depicts the mean percent parasite survival ± SEM (n = 4). Values represented by bars are the mean $IC_{50}$ or $IC_{90}$ ± standard error of the mean (SEM) (n = 4). Statistical significance was determined using unpaired t-tests (*$P < 0.05$, ***$P < 0.0005$, ****$P < 0.00005$) and is annotated above bars. The numbers below the bars indicate the $IC_{50}$ fold shifts between parental lines.

| Gene name | Gene ID | Full Gene Name | Associated Antimalarial Resistance | PMID | NF54 Genotype | PAT-039 Genotype |
|---|---|---|---|---|---|---|
| ubp1 | PF3D7_0104300 | ubiquitin carboxyl-terminal hydrolase 1 | Artemisinin | 31636063 | T116, S149 | T116, S149**F** |
| mrp1 | PF3D7_0112200 | multidrug resistance-associated protein 1 | Sulfadoxine-Pyrimethamine, Artemether-Lumefantrine | 19364873, 19807279 | I876 | I876**V** |
| acs8 | PF3D7_0215300 | acyl-CoA synthetase | – | – | E750 | E750**Q** |
| abci3 | PF3D7_0319700 | ABC transporter I family member 1 | Imidazopyridine, Carboxamide | 34233174 | K1973 | K1973**N** |
| carl | PF3D7_0321900 | cyclic amine resistance locus protein | Ganaplacide (KAF156) | 22096101 | K903 | K903**E** |
| sec24b | PF3D7_0405100 | protein transport protein Sec24B | – | – | S184 | S184**N** |
| dhfr | PF3D7_0417200 | dihydrofolate reductase | Sulfadoxine-Pyrimethamine, Cycloguanil | 2904149, 2183222 | N51, C59, S108 | N51**I**, C59**R**, S108**N** |
| pi4k | PF3D7_0509800 | phosphatidylinositol 4-kinase beta | Imidazopyrazine, 2-aminopyrimidine | 24284631, 28446690 | N877, N878 | N877, N878**D** |
| aat1 | PF3D7_0629500 | amino acid transporter | Chloroquine | 37169919 | S258 | S258**L** |
| pare | PF3D7_0709700 | prodrug activation and resistance esterase | AN13762 | 37158739 | M261 | M261**I** |
| px1 | PF3D7_0720700 | phosphoinositide-binding protein | Artemether-Lumefantrine | 40766679 | M1701, D1705 | M1701**I**, D1705**N** |
| dhps | PF3D7_0810800 | dihydropteroate synthase | Sulfadoxine-Pyrimethamine | 7925353 | K540 | K540**E** |
| psf3 | PF3D7_0914800 | GINS complex subunit | – | – | S61 | S61**N** |
| atg18 | PF3D7_1012900 | autophagy-related protein 18 | Artemisinin | 30367653 | V82 | V82**I** |
| act | PF3D7_1036800 | acetyl-CoA transporter 1 | Ganaplacide | 27642791 | T459 | T459**I** |
| gexp02 | PF3D7_1102500 | gametocyte exported protein 2 | – | – | S411 | S411**T** |
| fp2b | PF3D7_1115300 | cysteine proteinase falcipain 2b | Artemisinin | 21709259 | V147, H150, V157, T165, I167 | V147**F**, H150**N**, V175**A**, T165**M**, I167 |
| kic5 | PF3D7_1138700 | kelch13 interacting candidate 5 protein | Artemisinin | 36624300 | E497 | E497**Q** |
| atp4 | PF3D7_1211900 | non-SERCA-type Ca2+ -transporting P-ATPase | Cipargamin | 20813948 | T507 | T507**S** |
| ap2mu | PF3D7_1218300 | AP-2 complex subunit mu | Artemisinin, Quinine | 25691625 | F437 | F437**L** |
| mrp2 | PF3D7_1229100 | multidrug resistance-associated protein 2 | Mefloquine, Chloroquine, Quinine | 24372851 | S1527, L1531 | S1527**T**, L1531**I** |
| coronin | PF3D7_1251200 | coronin | Artemisinin | 30420498 | S183, K212 | S183**G**, K212 |
| k13 | PF3D7_1343700 | kelch protein | Artemisinin | 24352242 | C469 | C469**Y** |
| mdr2 | PF3D7_1447900 | multidrug resistance protein 2 | Pyrimethamine | 22314533 | S208, N229 | S208**N**, N229**D** |
| crt | PF3D7_0709000 | chloroquine resistance transporter | Chloroquine, Amodiaquine, Piperaquine, Quinine | 11090624, 19884511, 30115924,17163969 | CVMNK | CVMNK |
| mdr1 | PF3D7_0523000 | multidrug resistance protein 1 | Mefloquine, Amodiaquine, Lumefantrine, Quinine | 8426608, 27189525, 10706290 | N86, Y184 | N86, Y184 |

**Extended Data Figure 2. Full list of genes in the 24-marker SNP genotyping panel, associated antimalarial resistance and parental genotype.** Noted for each target in the panel is the common gene name, gene ID, full gene name, name of the antimalarial associated with mutations in each gene, and the parental genotype. Highlighted in bold is the amino acid substitution occurring due to non-synonymous mutations in PAT-039.

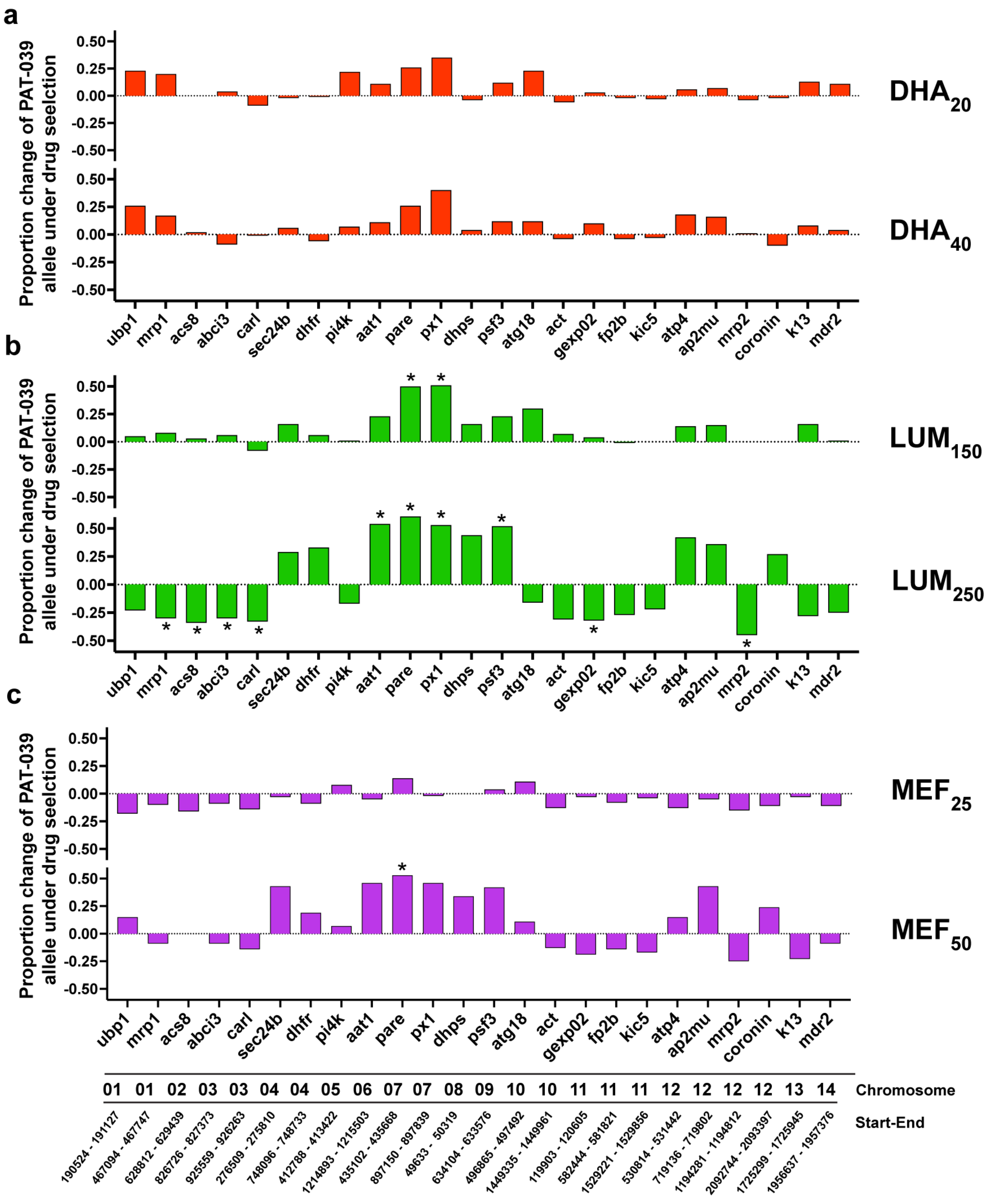


**Extended Data Figure 3. Concentration-dependent effects of dihydroartemisinin (DHA), lumefantrine (LUM) and mefloquine (MEF) selection on the proportion of PAT-039 alleles relative to controls in drug-selected progeny across 24 marker genotyping panel.** **a**, 20 nM and 40 nM DHA. **b**, 150 nM and 250 nM LUM. **c**, 25 nM and 50 nM MEF. Each bar represents the proportion of PAT-039 allele detected in drug-selected samples relative to that in untreated controls. Bars are colored according to each drug treatment condition. Statistical significance was determined using Z-score (*P < 0.05) and is annotated above bars. Annotated below marker names at the bottom of the figure are the chromosomal locations of each marker and the genomic position of the PCR amplicon used for nanopore sequencing and SNP genotyping. For the full list of marker gene names and IDs, refer to Extended Data Fig. 2.

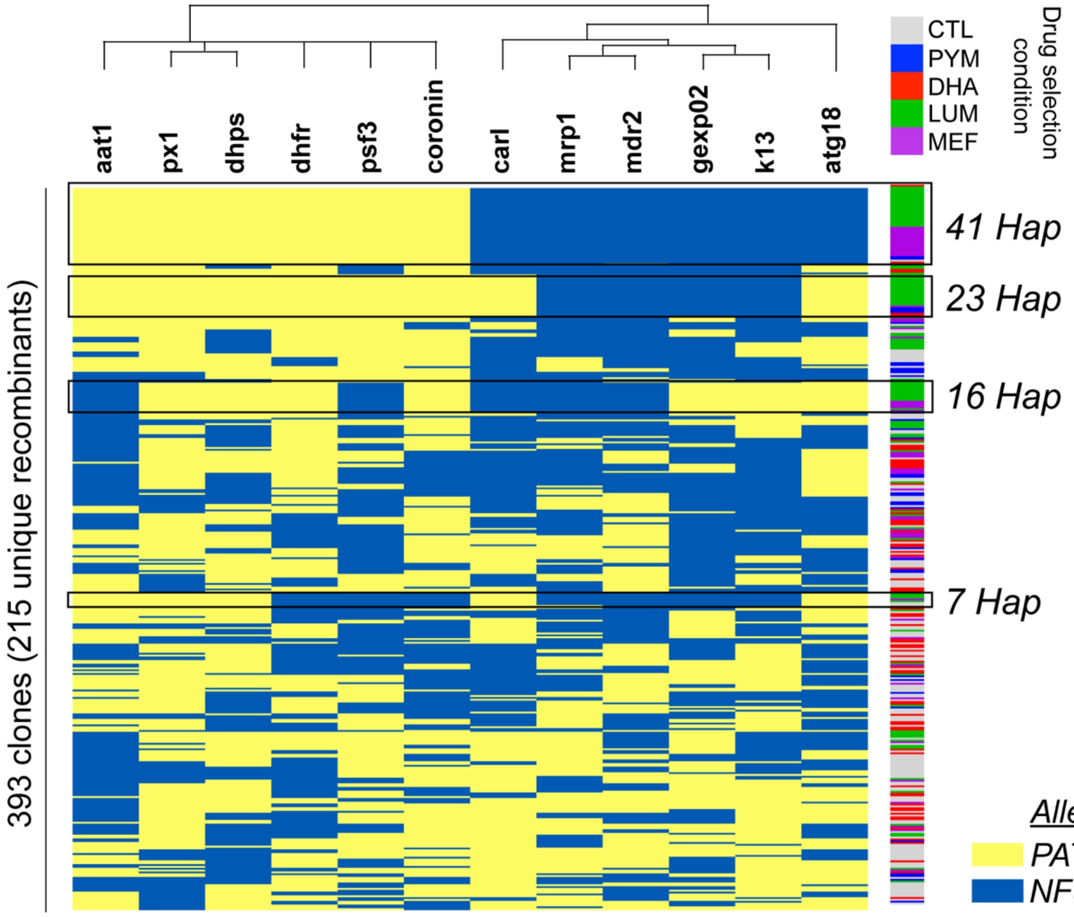


**Extended Data Figure 4. Allelic map of progeny clones arranged according to genotypic group.** Allelic map of progeny clones showing the parental allele inheritance among each of the 12 genes targeted using SNP marker panel across all 393 progeny clones genotyped. Alleles are colored according to the parental origin (NF54 or PAT-039). Color bar to the right of allelic map displays the drug-selection or control condition from which each clone is derived. Annotated are the four largest genotypic groups and the number of progeny constituting each group. For the full list of marker gene names and IDs, refer to Extended Data Fig. 2.

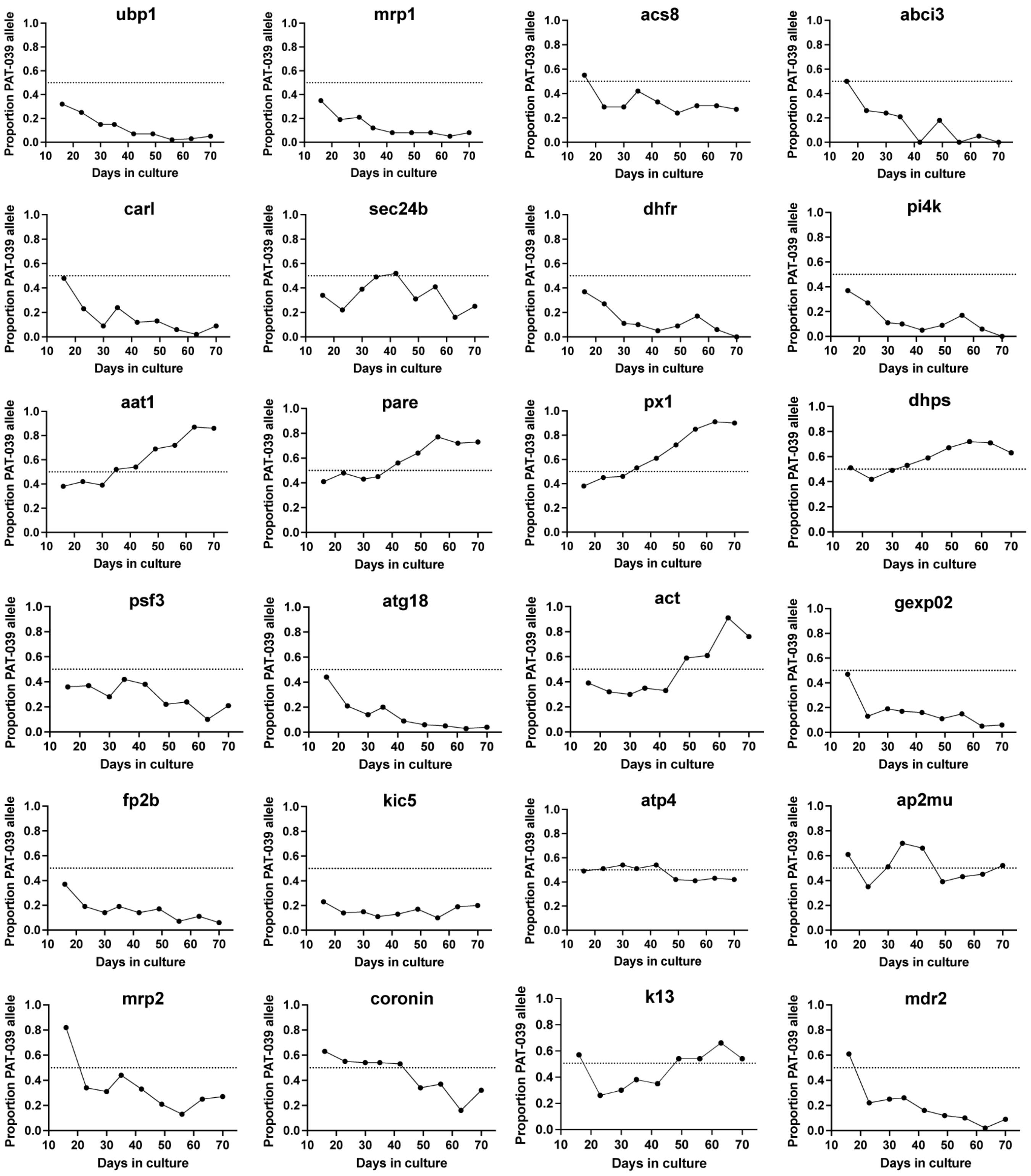


**Extended Data Figure 5. PAT-039 allele frequency measured in progeny pools in fitness assays across the 24-marker genotyping panel.** Each graph depicts the proportion of PAT-039 allele recorded at nine timepoints across 70 days in culture for each marker gene. Graphs are arranged in ascending chromosomal order for each marker. *Px1* is the marker associated with the highest parasite growth rate. For the full list of marker gene names and IDs, refer to Extended Data Fig. 2.

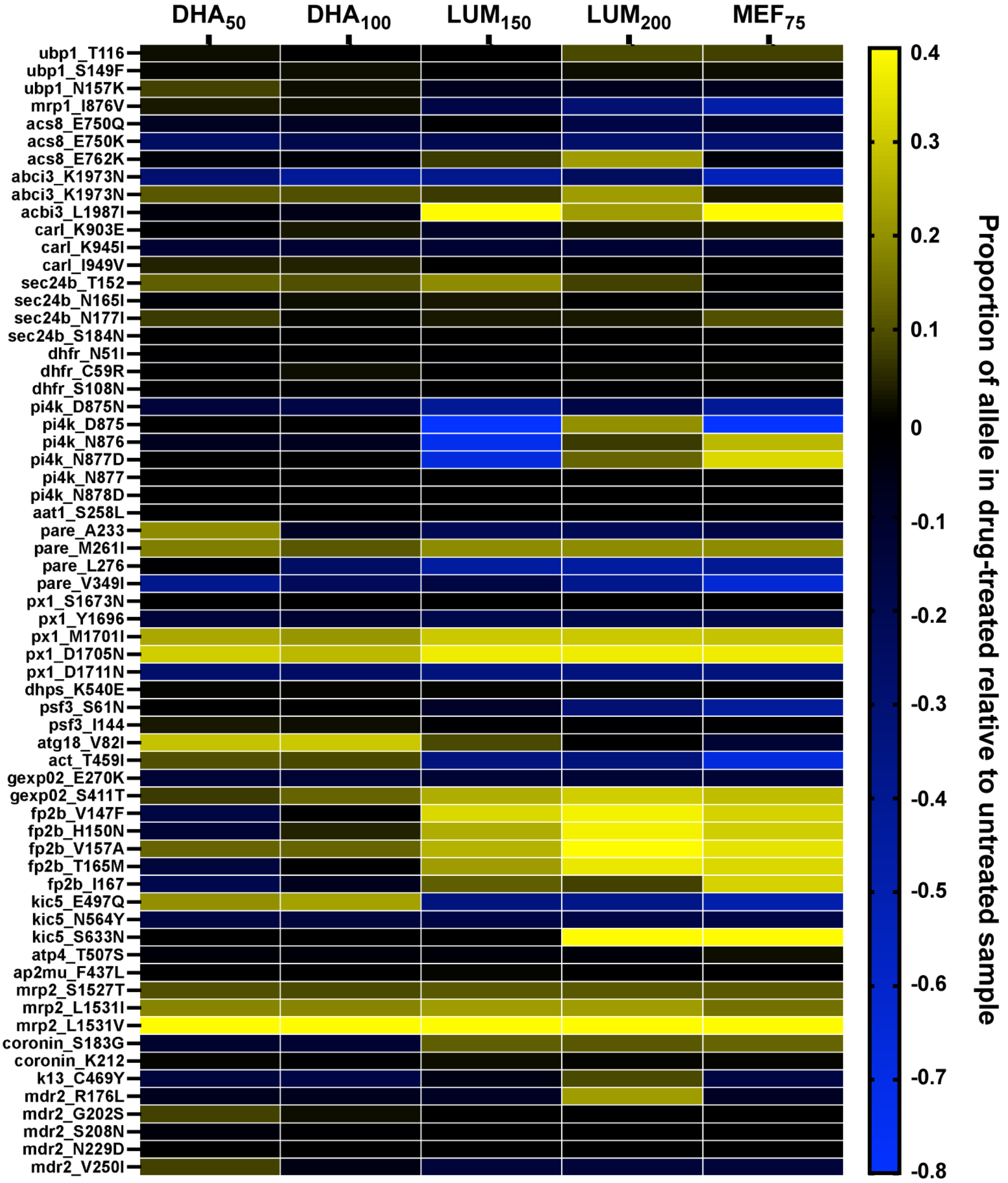


**Extended Data Figure 6. Full heat map displaying the change in the proportion of non-NF54 alleles in pooled Ugandan isolates under drug selection relative to controls for all SNPs in the 24-marker genotyping panel.** Individual cells represent the allele proportion for each marker gene SNP in each drug selection condition relative to untreated controls. The yellow-blue scale bar represents the change in the proportion of the allele in the drug-treated samples relative to the untreated control. SNPs are arranged in ascending chromosomal order. For the full list of marker gene names and IDs, refer to Extended Data Fig. 2.

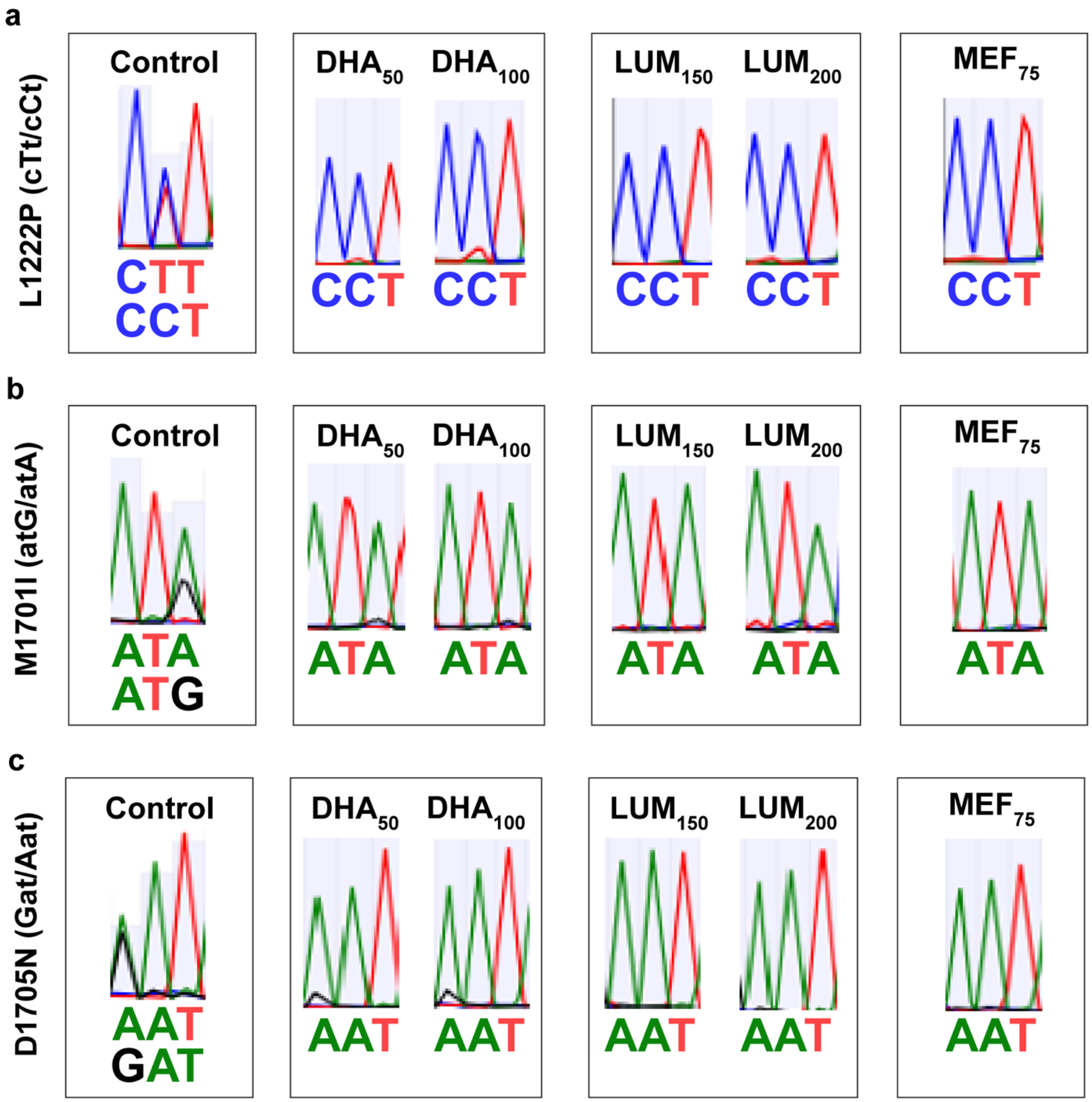


**Extended Data Figure 7. Genotyping of *px1* in the pooled Ugandan clinical isolates under different drug selection and control conditions determined by Sanger sequencing shows an enrichment for the PIN haplotype. a**, Representative Sanger sequencing chromatograms for sequencing of the pooled Ugandan clinical isolates at the px1 locus encoding L1222 in control and drug-selected samples. **b**, Sequencing at the locus encoding M1701 and (**c**) D1705. Annotated for each chromatogram is the codon at the relevant amino acid position. If the chromatogram indicates a mixed population at this position, both codons present are annotated.
