## Supplementary Figures & Tables for "A *Plasmodium falciparum* PX1 haplotype is associated with reduced susceptibility to artemisinin and lumefantrine"

### Supplementary Figures & Supplementary Tables

This file contains Supplementary Figures 1-5 & Supplementary Tables 1-5

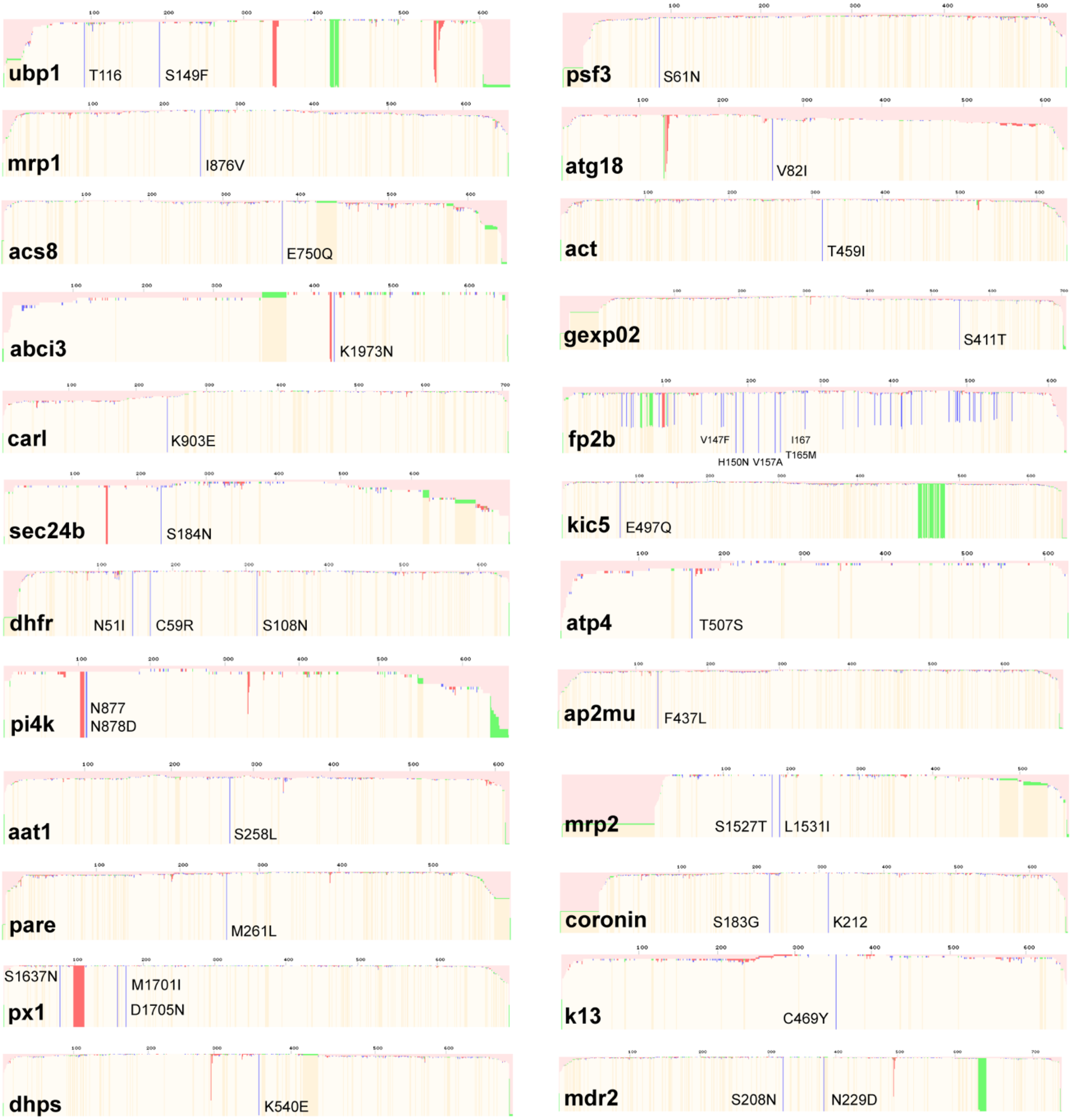


**Supplementary Figure 1. Representative images of SNP marker genotyping output for cross parent PAT-039.** Images depicting the nanopore sequence alignment for the PAT-039 line relative to the 3D7 reference for each marker gene in the 24-marker SNP genotyping panel. Annotated are the name of each marker gene and the SNPs present within the PCR amplicon. Markers are arranged in ascending chromosomal order. For the full list of marker gene names and IDs, refer to Extended Data Fig. 2.

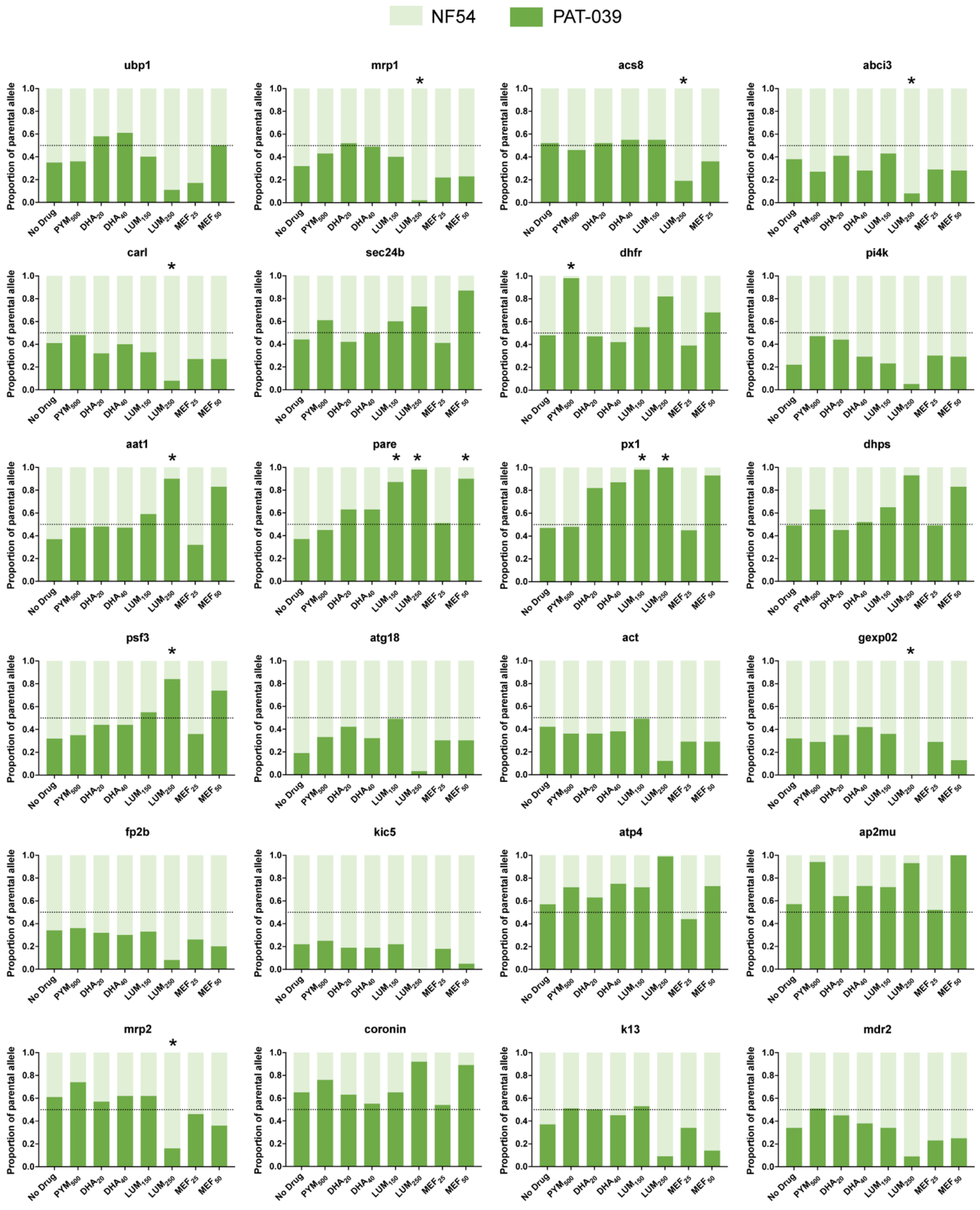


**Supplementary Figure 2. Proportion of each parental allele in drug-selected progeny pools and controls across the 24 marker SNP genotyping panel.** Graphs depict the proportion of PAT-039 and NF54 parental alleles recorded for every marker in the 24-marker genotyping panel in each drug selection condition applied on progeny pools and untreated controls. Each graph represents a different marker gene in the panel, arranged in ascending chromosomal order. Bars are colored according to the legend above the figure. Statistical significance was determined using Z-score (*P < 0.05) and is annotated above bars. For the full list of marker gene names and IDs, refer to Extended Data Fig. 2.

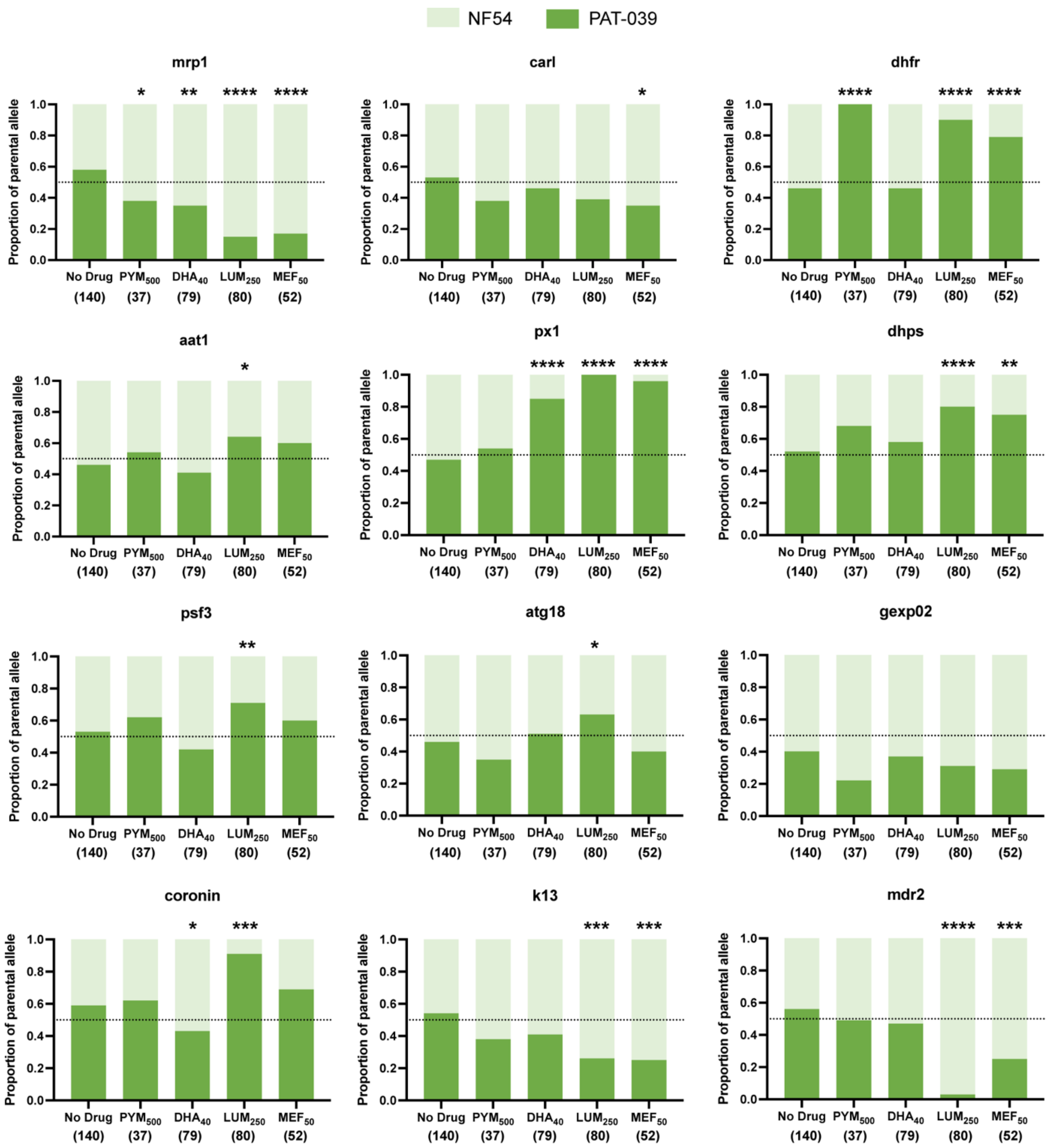


**Supplementary Figure 3. Proportion of each parental allele in progeny clones derived from drug selection and control conditions across the 12-marker SNP genotyping panel.** Graphs depict the proportion of clones with PAT-039 and NF54 parental alleles recorded for each marker in the 12-marker genotyping panel for clones derived from each drug-selection or control condition. Each graph represents a different marker gene in the panel, arranged in ascending chromosomal order. Bars are colored according to the legend above the figure. Statistical significance was determined using Fisher's Exact Test on raw counts (*$P < 0.05$, **$P < 0.005$, ***$P < 0.0005$, ****$P < 0.00005$) and is annotated above bars. For the full list of marker gene names and IDs, refer to Extended Data Fig. 2.

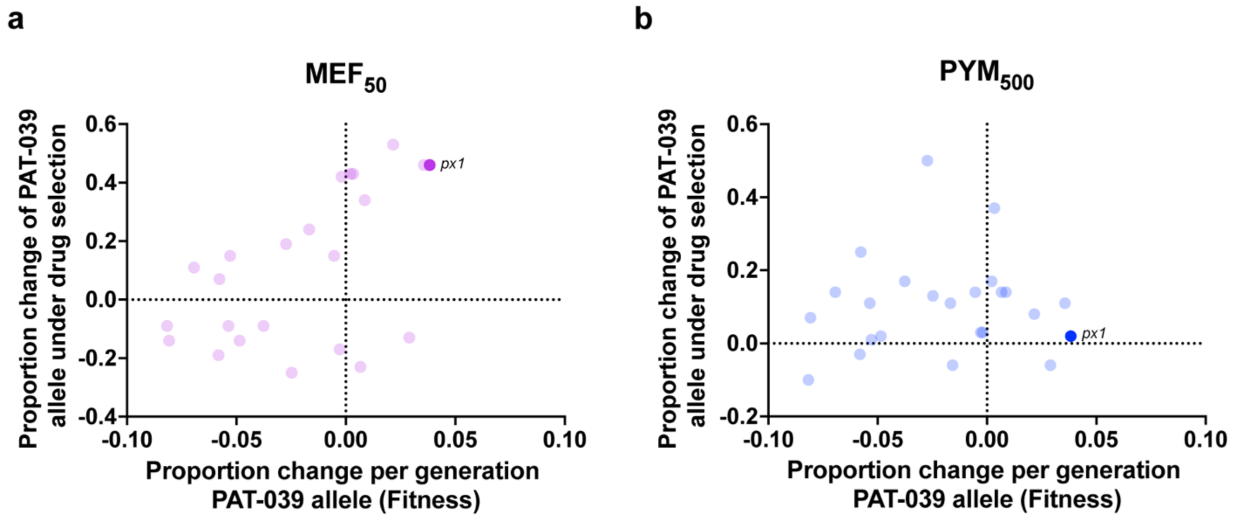


**Supplementary Figure 4. Fitness and drug-selection phenotype association for all markers under mefloquine (MEF) and pyrimethamine (PYM) selection.** Association between the fitness phenotype (data shown in Fig. 4b) and drug-selection phenotype (data shown in Fig. 2) for each marker gene. Each point represents an individual marker gene colored according to drug selection condition. Data points for *px1* are annotated.

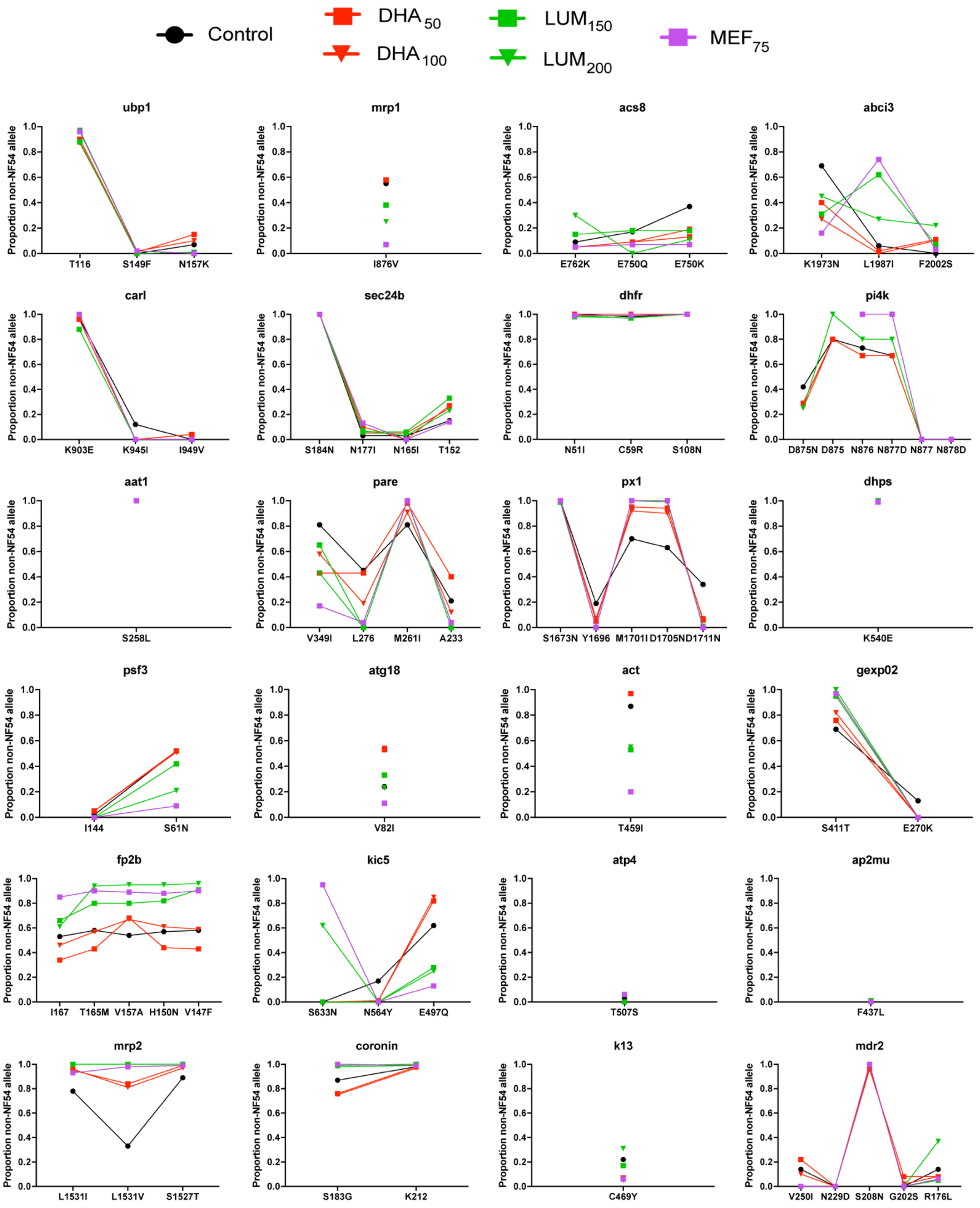


**Supplementary Figure 5. Non-NF54 allele frequency in pooled Ugandan isolates across all SNPs in the 24-marker genotyping panel.** Proportion of non-NF54 allele for every SNP encompassed by the 24-marker genotyping panel in each drug selection condition and control. Individual graphs depict the proportion of non-NF54 alleles across all SNPs for a distinct gene in the 24-marker genotyping panel and are arranged in ascending chromosomal order. Points and lines are colored according to the legend above the figure. For the full list of marker gene names and IDs, refer to Extended Data Fig. 2.

**Supplementary Table 1. Ring-stage survival (RSA), IC50 and Standard Error of Mean (SEM) values for dihydroartemisinin (DHA), lumefantrine (LUM) or mefloquine (MEF) for the cross parental lines NF54 and PAT-039 (n=4 independent experiments).**

| | | NF54 | PAT-039 | P-value |
|---|---|---|---|---|
| %RSA (72h) | Mean | 0.7 | 19.0 | 0.0001 |
| | SEM | 0.2 | 2.1 | |
| %RSA (Day5) | Mean | 1.3 | 49.0 | 0.0007 |
| | SEM | 0.4 | 7.5 | |
| DHA (AUC) | Mean | 1.2 | 24.0 | <0.0001 |
| | SEM | 0.2 | 2.3 | |
| DHA (16h) IC50 | Mean | 1.6 | 3.1 | 0.0004 |
| | SEM | 0.2 | 0.0 | |
| DHA (72h) IC50 | Mean | 1.5 | 2.5 | 0.0353 |
| | SEM | 0.1 | 0.3 | |
| LUM (72h) IC50 | Mean | 41.4 | 127.0 | <0.0001 |
| | SEM | 3.5 | 1.9 | |
| LUM (72h) IC90 | Mean | 121.0 | 260.0 | 0.0001 |
| | SEM | 4.8 | 14.7 | |
| MEF (72h) IC50 | Mean | 21.2 | 54.4 | <0.0001 |
| | SEM | 2.3 | 1.3 | |
| MEF (72h) IC90 | Mean | 38.2 | 106.0 | <0.0001 |
| | SEM | 2.7 | 6.2 | |

**Supplementary Table 2. Primer sequences and genomic positions for SNP genotyping markers.**

| Marker Name | Gene ID | Chromosome | Start | End | Forward Sequence | Reverse Sequence | SNPs Encompassed in PAT-039 |
|---|---|---|---|---|---|---|---|
| ubp1 | PF3D7_0104300 | Pf3D7_01_v3 | 190524 | 191127 | GTCACAACAAACCACCCGAGTC | CACTATTCTTCGTATCGGAGTCGTATG | T116, S149F |
| mrp1 | PF3D7_0112200 | Pf3D7_01_v3 | 467094 | 467747 | GACAAAAAGTAAGAATATGTTTAGCTAGAGC | CGTATTCCCTTTTACGAAATAAGCTTCC | I876V |
| acs8 | PF3D7_0215300 | Pf3D7_02_v3 | 628812 | 629439 | GGTGATGATTCTTTAGATGAAGCCTTAGC | CACTGGTATGGATATATTATCCTGTGGAAGTG | E750Q |
| abci3 | PF3D7_0319700 | Pf3D7_03_v3 | 826726 | 827373 | CGTTGGAGACAAGAAATGATACGGATG | GACATGGAAATTTTCAAAGATTTTCCTATTCTC | K1973N |
| carl | PF3D7_0321900 | Pf3D7_03_v3 | 925559 | 926263 | GCCCAGCCTTTTATGAAATTATATGTGG | CGCACAAGGTGCATAAATGCATG | K903E |
| sec24b | PF3D7_0405100 | Pf3D7_04_v3 | 276509 | 275810 | GCTGCTAGACAAAAGAGTCAATATATGAACC | CTCACCTGACTCCTCATTATCTTCATCC | S184N |
| dhfr | PF3D7_0417200 | Pf3D7_04_v3 | 748096 | 748733 | ACAAGTCTGCGACGTTTTCGATATTTATG | AGTATATACATCGCTAACAGA | N51I, C59R, S108N |
| pi4k | PF3D7_0509800 | Pf3D7_05_v3 | 412788 | 413422 | GTTGCTGATTTGGATGAAGATATTAGTCAC | CGTCATTATCATCGTCATTATCATCGTC | N877, N878D |
| aat1 | PF3D7_0629500 | Pf3D7_06_v3 | 1214893 | 1215503 | CTGGTGGTGTTAGATCTAGTACGGTAC | CATGCATTTGGTTGTTGAGAGAAGG | S258L |
| pare | PF3D7_0709700 | Pf3D7_07_v3 | 435102 | 435668 | GGGATTATCTATGGGGGGTAATGTTG | CTTGTTTGGGGGTATGGACAGC | M261I |
| px1 | PF3D7_0720700 | Pf3D7_07_v3 | 897150 | 897839 | GGACCACCACATGCAAACAC | GTTCTACCGGTACTGAATCTTCATTTTG | S1637N, M1701I, D1705N |
| dhps | PF3D7_0810800 | Pf3D7_08_v3 | 49633 | 50319 | TTGAAATGATAAATGAAGGTGCTAGT | CCAATTGTGTGATTTGTCCA | K540E |
| psf3 | PF3D7_0914800 | Pf3D7_09_v3 | 634104 | 633576 | GATTTAGTTGAAATACCATGTGTACCTG | GCATTTCTCCTTTTCGTAAAAATGAAATAAGG | S61N |
| atg18 | PF3D7_1012900 | Pf3D7_10_v3 | 496865 | 497492 | GGTTTCAAGATATACAACACGAACCC | CATCCAATAGAATTATCATGGGCATATATACTC | V82I |
| act | PF3D7_1036800 | Pf3D7_10_v3 | 1449335 | 1449961 | CATTGTACGAATTATTATTTCTACCCCCAATG | CATATATGTCCCTCCTATTTTAGGGTCTG | T459I |
| gexp02 | PF3D7_1102500 | Pf3D7_11_v3 | 119903 | 120605 | CTCAAGTATGAAAACTTTGAAGAGTATGC | CCACATATCCCTTAAGGTATTCACATC | S411T |
| fp2b | PF3D7_1115300 | Pf3D7_11_v3 | 582444 | 581821 | CCCTAATGGCAAGAAATTTATCGTCTC | CTACGGAACCTATACTACTAAAGGCCC | V147F, H150N, V157A, T165M, I167 |
| kic5 | PF3D7_1138700 | Pf3D7_11_v3 | 1529221 | 1529856 | GCAAATCGTTATCAACAGGTACCAAG | CCTTTGTTTATATTTCCAACGGTAGGC | E497Q |
| atp4 | PF3D7_1211900 | Pf3D7_12_v3 | 530814 | 531442 | CCGGTACATTAACTGAAGGAAAAATGAC | CTTCTATCAAGTAATCTATCAGGTGCACC | T507S |
| ap2mu | PF3D7_1218300 | Pf3D7_12_v3 | 719136 | 719802 | CTTCACACCACCAGATGGTACATTTG | CCCGATTCTGTTAAATACTTGATCCAC | F437L |
| mrp2 | PF3D7_1229100 | Pf3D7_12_v3 | 1194281 | 1194812 | GTTAATAGCACATGGTATTGTTAAATCGGC | GTTAGCGCCAGAACAATTAATTGTACG | S1527T, L1531I |
| coronin | PF3D7_1251200 | Pf3D7_12_v3 | 2092744 | 2093397 | GATGAGAATGTGAATGAGGTAAAGGATCC | CATTTGTATACGTCACACATACGTTTAGG | S183G, K212 |
| k13 | PF3D7_1343700 | Pf3D7_13_v3 | 1725299 | 1725945 | GATTGATATTAATGTTGGTGGAGC | TACCCATGCTTTCATACGATGAT | C469Y |
| mdr2 | PF3D7_1447900 | Pf3D7_14_v3 | 1956637 | 1957376 | GGTGTATGTATTATCCAGACATATAACCGG | GCTTGGCCACAAATATGGTAAAAG | S208N, N229D |

**Supplementary Table 3. Mean, SEM and P-values for percentage change per generation of PAT-039 allele in progeny fitness assays.**

| Marker Name | Mean | SEM | n | P-Value |
|---|---|---|---|---|
| ubp1 | -0.0529 | 0.05528 | 8 | 0.3706 |
| mrp1 | -0.0536 | 0.04003 | 8 | 0.2223 |
| acs8 | -0.0158 | 0.03557 | 8 | 0.6713 |
| abci3 | -0.0817 | 0.04495 | 3 | 0.2109 |
| carl | -0.0807 | 0.1008 | 7 | 0.9572 |
| sec24b | 0.00213 | 0.05153 | 8 | 0.9683 |
| dhfr | -0.0273 | 0.03178 | 7 | 0.874 |
| pi4k | -0.0577 | 0.07038 | 7 | 0.4436 |
| aat1 | 0.03563 | 0.01752 | 8 | 0.0816 |
| pare | 0.02163 | 0.01232 | 8 | 0.1226 |
| px1 | 0.03829 | 0.006435 | 7 | 0.001 |
| dhps | 0.00863 | 0.01332 | 8 | 0.5379 |
| psf3 | -0.002 | 0.05611 | 8 | 0.9726 |
| atg18 | -0.0694 | 0.04588 | 8 | 0.1743 |
| act | 0.029 | 0.02982 | 8 | 0.3632 |
| gexp02 | -0.0581 | 0.05655 | 8 | 0.3383 |
| fp2b | -0.0485 | 0.04767 | 8 | 0.3428 |
| kic5 | -0.0029 | 0.04087 | 8 | 0.9459 |
| atp4 | -0.0054 | 0.01023 | 8 | 0.6156 |
| ap2mu | 0.00338 | 0.03697 | 8 | 0.9298 |
| mrp2 | -0.0248 | 0.05126 | 8 | 0.6439 |
| coronin | -0.0168 | 0.0452 | 8 | 0.7219 |
| k13 | 0.00663 | 0.03587 | 8 | 0.8587 |
| mdr2 | -0.0376 | 0.09382 | 8 | 0.7004 |

**Supplementary Table 4. Sample collection and genotypes of drug-resistance markers for the 18 Ugandan isolates that constitute the pool.**

| | | | Genotypes in Drug Resistance-Associated Markers | | | | | |
|---|---|---|---|---|---|---|---|---|
| Parasite line | Year collected | Field site | *k13* [1] | *px1* [2] | *mdr1* [3] | *crt* [4] | *dhfr* [5] | *dhps* [6] |
| PAT-107 | 2022 | Patongo, Agago (North Uganda) | K189T | LIDN | NYD | CVMNK | IRNI | GK[#] |
| PAT-070 | 2022 | Patongo, Agago (North Uganda) | K189T | PIND | NFD | CVMNK | IRNI | GE |
| PAT-088 | 2022 | Patongo, Agago (North Uganda) | C469Y[#] | PIND | NFD | CVMNK | IRNI | GE |
| PAT-148 | 2022 | Patongo, Agago (North Uganda) | WT | PIND | NYD | CVMNK | IRNI | GE |
| PAT-154 | 2022 | Patongo, Agago (North Uganda) | A675V[#] | PIND | NYD | CVMNK | IRNI | GE |
| PAT-171 | 2022 | Patongo, Agago (North Uganda) | K189T | PIND | NYD | CVMNK | IRNI | GE |
| BUS-111 | 2022 | Busiu, Mbale (East Uganda) | C469Y | L[#]MDD[#] | NYD | CVMNK | IRNI | GE |
| BUS-140 | 2022 | Busiu, Mbale (East Uganda) | A627V | PIND | NFD | CVMNK | IRNI | GE |
| BUS-122 | 2022 | Busiu, Mbale (East Uganda) | K189N | PIND | NFD | CVMNK | IRNI | GE |
| U03-299 | 2023 | Tororo (East Uganda) | K189T | L[#]M[#]D[#]D[#] | NFD | CVMNK | IRNI | GE |
| U03-193 | 2023 | Tororo (East Uganda) | WT | LI[#]N[#]N[#] | NFD | CVMNK | IRNL[#] | GE |
| U03-087 | 2022 | Tororo (East Uganda) | K189T | LMDD[#] | NYD | CVMNK | IRNI | GE |
| U03-150 | 2023 | Tororo (East Uganda) | WT | LMDN | NFD | CVMNK | IRNI | GE |
| U03-208 | 2023 | Tororo (East Uganda) | N217H | LMDN | NYD | CVMNK | IRNI | GE |
| U03-140 | 2023 | Tororo (East Uganda) | WT | PIND | NYD | CVMNK | IRNI | GE |
| U03-181 | 2023 | Tororo (East Uganda) | WT | PI[#]N[#]D | NYD | CVMNK | IRNI | GE |
| U03-242 | 2023 | Tororo (East Uganda) | WT | P[#]IN[#]D | NYD | CVMNK | IRNI | GE |
| U03-300 | 2023 | Tororo (East Uganda) | WT | PID[#]D | NYD | CVMNK | IRNL | GE |

[1] *k13* genotype across full-length gene

[2] *px1* genotype at positions L1222P, M1701I, D1705N, D1711N; Wild-type: LMDD

[3] *pfmdr1* genotype at positions N86Y, Y184F, D1246Y; Wild-type: NYD

[4] *pfcrt* genotype at positions 72-76; Wild-type: CVMNK

[5] *dhfr* genotype at positions N51I, C59R, S108N, I164L; Wild-type: NCSI

[6] *dhps* genotype at positions at A437G, K540E; Wild-type: AK

[#] mixed genotypes (40-70%) with major genotype shown at that position

**Supplementary Table 5. Genotypes of drug-resistance markers and phenotypes depicted by Ring-Stage Survival (RSA) and IC50s mean values and SEM values of Ugandan clinical isolates that were profiled in drug susceptibility assays and studied herein.**

| | Genotypes in Drug Resistance-Associated Markers | | | %RSA | | | Lumefantrine IC50 | | | Mefloquine IC50 | | |
|---|---|---|---|---|---|---|---|---|---|---|---|---|
| Parasite | *k13* | *mdr1* | *px1* | Mean | SEM | n | Mean | SEM | n | Mean | SEM | n |
| PAT-070 | K189T | Y184F | PIN | 0.3 | 1.0 | 3 | 65.0 | 32.8 | 2 | 42.8 | 16.1 | 2 |
| PAT-171 | K189T | WT | PIN | -1.5 | 0.6 | 4 | 60.5 | 2.2 | 2 | 36.0 | 13.3 | 2 |
| DHA50_8 | K189T | Y184F | PIN | 2.8 | 0.6 | 2 | 67.2 | 5.6 | 2 | 60.0 | 5.0 | 2 |
| DHA100_1 | K189T | WT | PIN | 2.2 | 0.2 | 4 | 54.4 | 4.8 | 4 | 33.2 | 2.4 | 4 |
| DHA100_3 | K189T | WT | PIN | 18.2 | 10.5 | 5 | 72.2 | 9.4 | 5 | 34.4 | 2.6 | 5 |
| DHA100_6 | K189T | WT | PIN | 3.9 | 1.1 | 4 | 71.7 | 9.4 | 4 | 35.2 | 3.8 | 4 |
| BUS-122 | K189N | Y184F | PIN | 4.9 | 0.9 | 4 | 71.4 | 13.3 | 4 | 47.8 | 6.3 | 4 |
| PAT-149 | K189T | Y184F | PIN | 0.7 | 0.4 | 2 | 51.8 | 0.6 | 2 | 31.3 | 1.3 | 2 |
| U03-087 | K189T | WT | LMD | 0.9 | 1.5 | 3 | 32.1 | 8.8 | 3 | 27.5 | 0.5 | 3 |
| PAT-106 | WT | WT | PIN | 5.8 | 8.1 | 3 | 79.5 | 8.7 | 3 | 37.8 | 2.5 | 3 |
| U03-140 | WT | WT | PIN | 0.7 | 0.9 | 3 | 63.5 | 8.2 | 2 | 42.3 | 3.6 | 2 |
| DHA50_2 | WT | WT | PIN | 0.5 | 0.2 | 3 | 71.5 | 9.1 | 3 | 43.8 | 5.1 | 3 |
| DHA50_3 | WT | WT | PIN | 0.9 | 0.9 | 3 | 61.0 | 14.3 | 3 | 30.5 | 2.9 | 3 |
| DHA50_4 | WT | T199S | PIN | 0.6 | 0.4 | 3 | 34.7 | 6.7 | 3 | 27.3 | 4.8 | 3 |
| DHA100_8 | WT | WT | PIN | 18.5 | 6.8 | 5 | 128.0 | 16.7 | 4 | 61.9 | 5.8 | 4 |
| DHA100_4 | WT | WT | PIN | 4.6 | 1.5 | 4 | 50.2 | 9.8 | 4 | 20.5 | 1.8 | 4 |
| DHA100_5 | WT | Y184F | PIN | 2.5 | 1.1 | 2 | 44.9 | 0.0 | 2 | 35.7 | 2.9 | 2 |
| PAT-148 | WT | WT | PIN | 17.0 | 4.1 | 4 | 129.0 | 9.7 | 3 | 61.5 | 2.8 | 3 |
| PAT-023 | WT | Y184F | PIN | 4.9 | 0.9 | 3 | 173.0 | 25.5 | 3 | 91.1 | 16.1 | 3 |
| U03-150 | WT | Y184F | LMD | -0.4 | 2.2 | 3 | 30.1 | 8.6 | 2 | 26.0 | 6.7 | 2 |
| PAT-039 | C469Y | WT | PIN | 19.0 | 2.1 | 4 | 114.0 | 11.4 | 3 | 52.3 | 2.6 | 3 |
| HL2210 | A675V | Y184F | PIN | 2.1 | 0.4 | 5 | 50.3 | 9.6 | 4 | 22.6 | 5.2 | 4 |
| U03-208 | N217H | WT | LMD | 1.9 | 2.5 | 4 | 34.1 | 23.6 | 2 | 26.6 | 1.4 | 2 |
| HL2208 | T348I | Y184F | PIN | 1.1 | 0.4 | 3 | 67.2 | 19.5 | 3 | 35.2 | 9.4 | 3 |
| BUS-030 | T348I | Y184F | LMD | -0.5 | 1.2 | 3 | 52.1 | 7.2 | 3 | 28.4 | 2.7 | 3 |
| BUS-056 | T348I | Y184F | LMD | 0.9 | 1.0 | 2 | 42.1 | - | 1 | 26.5 | - | 1 |
| TDH-144 | T348I | Y184F | LMD | 2.9 | 2.2 | 2 | 60.6 | 7.3 | 2 | 26.7 | 2.5 | 2 |
| TDH-191 | A675V | WT | LMD | 6.1 | 1.1 | 4 | 59.8 | 12.4 | 3 | 51.1 | 4.4 | 3 |
| PAT-045 | A675V | Y184F | PIN | 3.9 | 1.4 | 4 | 95.3 | 14.7 | 4 | 76.9 | 0.6 | 3 |
| PAT-046 | C469Y | WT | LMD | 1.5 | 0.6 | 4 | 41.1 | 7.6 | 3 | 46.6 | 3.0 | 3 |
| BUS-025 | C469Y | Y184F | PIN | 11.3 | 3.1 | 4 | 95.8 | 14.5 | 3 | 63.4 | 6.3 | 3 |
| PAT-026 | C469Y | Y184F | PIN | 7.8 | 1.3 | 4 | 59.8 | 9.3 | 3 | 55.1 | 2.6 | 3 |
| PAT-038 | WT | Y184F | PIN | 1.2 | 0.7 | 3 | 82.1 | 15.2 | 4 | 22.5 | 2.2 | 4 |
| BUS-041 | K189T/K189 | Y184F | PIN | 6.6 | 1.0 | 3 | 38.1 | 9.3 | 3 | 29.2 | 4.3 | 3 |
| BUS-035 | A621/T149S | Y184F | PIN | 2.2 | 1.8 | 4 | 98.8 | 10.6 | 3 | 52.0 | 2.8 | 3 |
| PAT-015 | WT | WT | PIN | 2.3 | - | 1 | 130.5 | 27.9 | 4 | 46.5 | 4.7 | 4 |
